# Manipulation of *in planta* ethylene levels modulates the metabolome of *Populus tremula x tremuloides* in a microbial-dependent manner

**DOI:** 10.1101/2025.03.17.643694

**Authors:** F. Fracchia, F. Guinet, S. Warion, M. Bednareck, C. Chervin, N. Engle, T.J. Tschaplinski, C. Veneault-Fourrey, A. Deveau

**Author notes:** Corresponding authors: Aurélie Deveau,; Claire Veneault-Fourrey.

## Abstract

**Background:** The assembly of tree microbiota is a dynamic process that varies across space and time, influenced by both biotic and abiotic factors. Plants have developed defence strategies, mediated by phytohormones such as ethylene, to manage its interactions with pathogens and, in certain non-perennials, interactions with the commensal microbiota. However, whether and how ethylene regulation affects the assembly of tree microbiota is unknown.

**Results:** We investigated the assembly of fungal and bacterial communities in poplars (*P. tremula* x. *P. tremuloides* ‘T89’), altered either in ethylene biosynthesis (*ACO1*), or in ethylene perception (etr1.1), by combining high-throughput amplicon sequencing with confocal microscopy. In parallel, we characterised root exudates, as well as root and shoot metabolomes of the different poplar lines grown on sterile soil and in the presence of microbiota using GC-MS. Alteration of ethylene levels had little impact on the metabolome of sterile poplars, but led to differential primary and secondary metabolic responses in the root and shoots of poplars colonised by microorganisms. These metabolomic changes were associated with a decrease in fungal colonisation of shoots, particularly by saprophytes in the early stages, whereas reduced ethylene stimulated root colonisation by fungi. Conversely, arbuscular mycorrhizal fungal and bacterial communities, as well as root exudation, were little affected by changes in ethylene production.

**Conclusion:** The findings of this study suggest a potential dual role for ethylene in poplar, whereby its levels may either promote or inhibit microbial growth and activity, depending on the concentration, microbial trophic guild, and poplar organ.

## BACKGROUND

Plant microbiota, primarily bacteria and fungi, play a major role in plant health and development aiding water and nutrient uptake, and biotic and abiotic stress resistance. Microbial communities vary by plant compartments. In nature, soil represents the main reservoir of microorganisms for root colonization, under the influence of edaphic parameters and environmental factors (Uroz et al. 2016). In the rhizosphere, root exudates and volatiles, which are under the influence of the plant genotype, promote the first recruitment of microbial communities (Bonito et al. 2014; Cregger et al. 2018; Reinhold-Hurek et al. 2015; Sasse, Martinoia, and Northen 2018). Then, plant genotype and microbe-microbe interactions regulate the establishment of the endospheric microbiota, in both above and belowground compartments (Gottel et al. 2011; Hacquard and Schadt 2015; Lareen, Burton, and Schäfer 2016).

Reshaping of the microbial communities from the soil, rhizosphere, and the root could impact plant growth and its responses to environmental constraints, as well as stress, can lead to changes in microbial communities (Kiers et al. 2011; Timm et al. 2018; Vandenkoornhuyse et al. 2015).

Ethylene is a crucial plant hormone in regulating development (growth, senescence, leaf expansion, and root initiation) and stress response (Dubois, Van den Broeck, and Inzé 2018; Pattyn, Vaughan-Hirsch, and Van De Poel 2021). To maintain the balance between growth and defence, the level of ethylene or ethylene-signalling is tightly controlled. In brief, plant ethylene is synthesised from methionine via two main enzymes, the 1-aminocyclopropane-1-carboxylic acid (ACC) synthase (ACS) and ACC oxidase (ACO). Then ethylene is perceived by receptors (ETR) and triggers the signalling cascade resulting in the activation of ethylene transcription factors (ERF) and the ethylene mediated response (Dubois, Van den Broeck, and Inzé 2018). The plant-associated microbiota can significantly modulate host ethylene signalling. Some microbes increase ethylene levels by producing ACC oxidase (a microbial ethylene-forming enzyme), by stimulating ACC synthase activity in plants, or by indirectly affecting other plant hormones. Others decrease ethylene production by breaking down its precursor, ACC. Additionally, microbes that are either beneficial or pathogenic can modulate ethylene responses by producing hormones or effectors that interact with the ethylene signalling pathway (Arteca and Arteca 2008; Guan et al. 2015; Kloppholz, Kuhn, and Requena 2011). In addition to directly influencing ethylene levels, the microbiota plays a key role in shaping how plants perceive and respond to stress. Ravanbakhsh et al., (2018) suggest that ethylene signalling functions as part of a hologenome-level stress response, where genetic traits from both the plant and its associated microbiota are activated in response to environmental stressors. From this perspective, the modulation of ethylene can be the reflection of the co-evolution of plant and microbial traits, demonstrating their integrated adaptation to stress (Ravanbakhsh et al. 2018). Many studies have investigated the role of ethylene in shaping the microbiota, with inconsistent results. Some studies showed that *A. thaliana* mutants that are unable to sense ethylene (*EIN2*, *etr1*) or mutants that over-produce ethylene transcription factors, ERFs, exhibited an alteration of leaf bacterial communities, and root bacterial and fungal communities (Bodenhausen et al. 2014; Kudjordjie, Sapkota, and Nicolaisen 2021), while another experiment involving the ethylene-insensitive *ein2* and *etr1 A. thaliana* mutants did not show any significant shift in rhizosphere bacterial community composition (Doornbos et al. 2011). In a bi-partite interaction, Plett et al. (2014) observed an impaired *in planta* fungal growth of the ectomycorrhizal fungus (EMF) *Laccaria bicolor* in the roots of the transgenic poplar over-expressing the ACO1 enzyme. Fu et al. (2021) observed significant differences of rhizosphere bacterial communities in the Nr transgenic tomatoes (disrupted in the ethylene receptor ETR3), whereas French et al., (2019) detected alteration of both root and rhizosphere bacterial communities in ACD transgenic tomatoes (constitutively degrades ACC, the precursor to ethylene), but not in the Nr transgenic plants. Another interesting outcome from this study was to correlate the shift of microbial communities in Nr transgenic lines with changes in root exudation (Fu et al. 2021). Nevertheless, findings are inconsistent and most of the previous works have focused on rhizosphere bacterial communities in annual plants. In woody plants, exogenous ethylene stimulates xylem secondary growth and inhibits shoot height growth (Felten et al. 2018; Love et al. 2009). However, no studies addressed the role of ethylene in shaping microbial communities in woody perennial plants, nor its effects on tree metabolism.

Earlier research on poplar demonstrated that colonization of poplar root systems by bacteria and fungi occurs as a highly dynamic and successional process, with early colonizers-fast-growing opportunistic species-being replaced over time by specialized species like mycorrhizal fungi and endophytes (Fracchia et al. 2021). Similarly, (Dove et al. 2021) observed over five months that poplar roots showed increasing colonization by ectomycorrhizal fungi alongside reductions in saprotroph and pathogen populations, suggesting a progression towards more stable and beneficial microbial communities. In this study, we asked whether the ethylene level in poplar trees impacts its microbiota. We tested two main hypotheses: the alteration of ethylene perception and biosynthesis (1) influences ethylene production, poplar development, exudates composition, as well as root and shoot metabolomes (2) impacts soil, rhizosphere as well as poplar above and belowground fungal and bacterial communities.

## RESULTS

To test these hypotheses, we planted transgenic poplars *P. tremula x P. tremuloides* T89 lines generated by Love et al. (2009), and described as either insensitive (35S::At*etr1-1* lines 1E and 3A further named Etr1.1-NR1E and Etr1.1-NR3A), or ethylene-overproducing through *PttACO1* (1-Aminocyclopropane-1-Carboxylic Acid Oxidase) constitutive expression (further named ACO1) in natural and microbe-free soils (sterilized with gamma-irradiation) and we assessed both microbial colonisation and metabolomic responses (**Fig. 1**).

**Fig 1.**
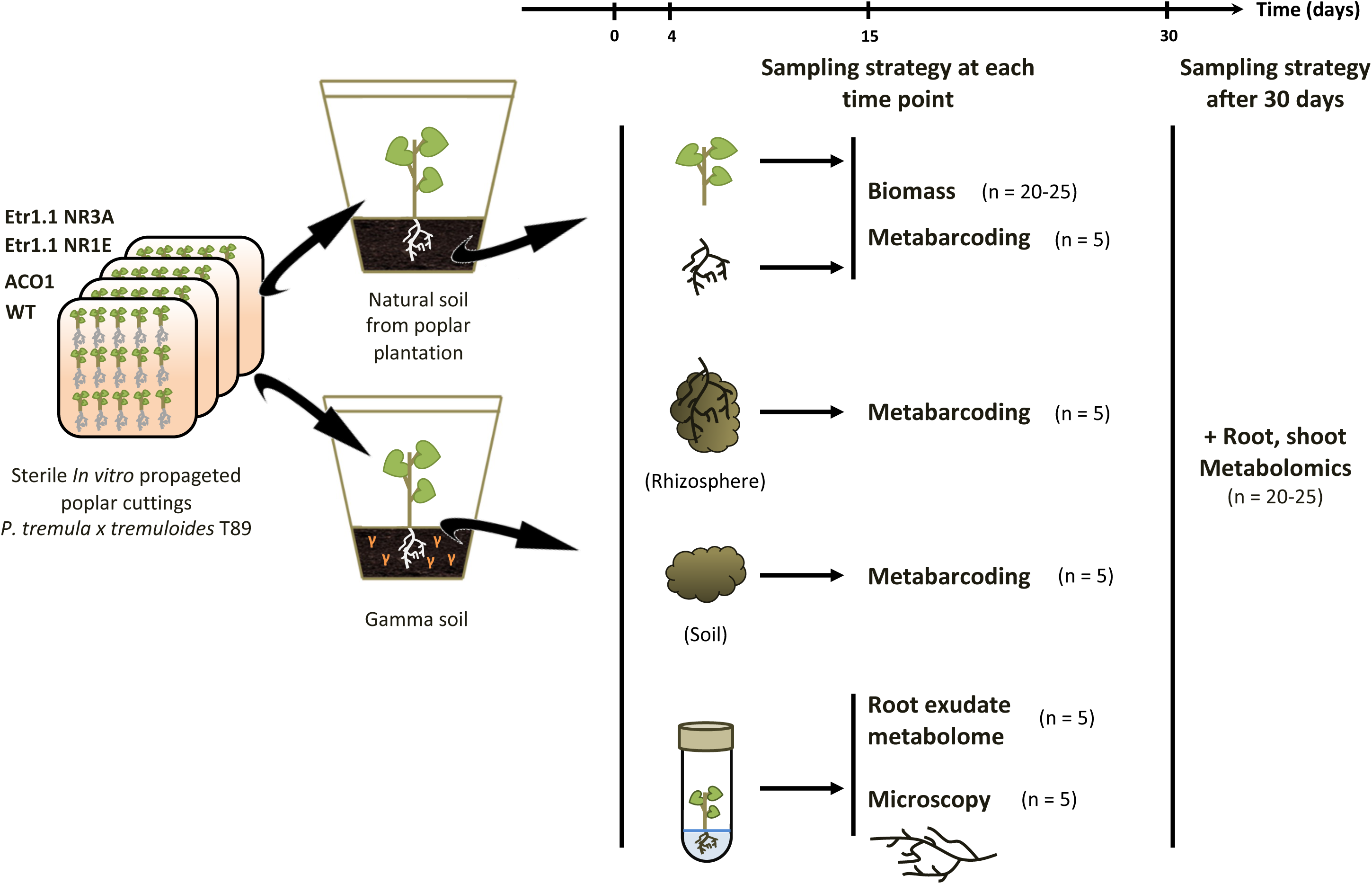
Schematic representation of the experimental design of the study. The seedlings of wild type poplar (T89) and transgenic lines grown in in vitro conditions were transplanted in microcosms containing either natural or gamma irradiated soils and grown over 30 days. At each time point in natural soil conditions, soil, rhizosphere, root, and shoot were sampled to assess the microbial colonisation. Over time, and in both soil types, we collected root exudates as well as root and shoot metabolomes.

### Alteration of *in planta* ethylene levels modifies root development without impairing metabolites in root exudates, roots, and shoots

Even though all transgenic lines over-expressed their transgenes after qPCR quantification (data not shown), we quantified the amount of ethylene produced by *in vitro* poplar transgenic lines either insensitive to ethylene (Etr1.1-NR1E and Etr1.1-NR3A) or overproducing the last enzyme involved in ethylene biosynthesis (ACO1) in comparison with wild type poplar by gas chromatography. Etr1.1-NR1E was the only transgenic line producing significantly 1.5x more ethylene in comparison with wild type T89 (WT) (LM, TukeyHSD post-hoc test, p.value ≤ 0.05), even though Etr1.1-NR3A displayed a tendency of ethylene overproduction (**Fig. 2A**). By contrast, the over-expression of ACO1 did not result in increased ethylene production, but instead tended to reduce ethylene production (**Fig. 2A**). Overall, insensitivity to ethylene triggered a positive feedback loop with increasing ethylene production, while the over-expression of ACO1 tended to result in a negative feedback loop and 25% decrease ethylene production in our conditions. Based on these results, we decided to keep T89 as WT-control, Etr1.1-NR1E as ethylene-overproducer, ACO1 as ethylene-low producer compared to Etr1.1-NR1E, and the transgenic line Etr1.1-NR3A was removed from further analyses.

**Fig. 2.**
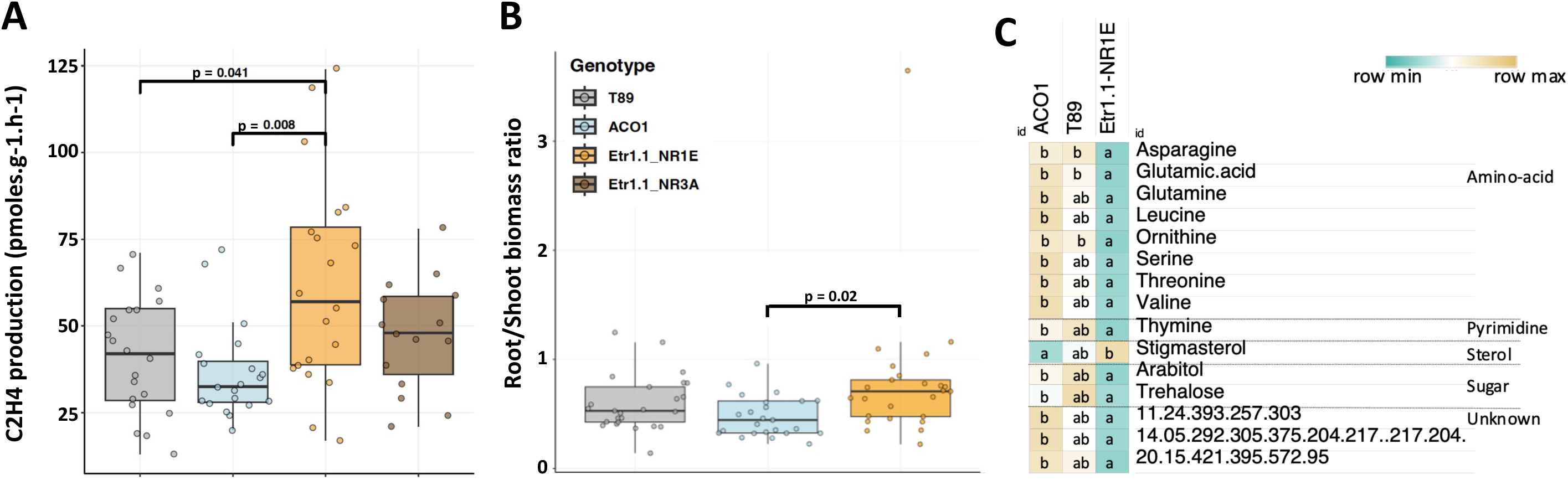
Ethylene production, fresh root and shoot biomass ratios, and metabolomes of poplar transgenic lines altered in ethylene perception/biosynthesis grown in sterile conditions. (A) Ethylene production of transgenic poplars and wild type (pmoles.g^-1^.h^-1^, LM, TukeyHSD post-hoc test, p value ≤ 0.05, n = 20). (B) Root/shoot fresh biomass ratio of transgenic and wild type poplars (LM, TukeyHSD post-hoc test, p value ≤ 0.05, n = 20-25). (C) List of root metabolites showing significant differences in abundance between the transgenic line ACO1, Etr1.1.NR1E, and wild type (Kruskall-Wallis with FDR corrections, Dunn post-hoc test, p adjusted ≤ 0.05, n = 20-25, Table S1A). Each row represents the z-score value of a metabolite in the three poplar genotypes. Unknown metabolites were named by retention time (min) followed by key mass-to-charge ratios (m/z) and putative metabolite classes are indicated.

Given that ethylene is involved in the regulation of developmental processes, we assessed the growth of the transgenic lines and of the T89 grown in the microcosms on sterile soil. After 30 days of growth, the root-shoot ratio increased significantly by 1.5x (LM, TukeyHSD post-hoc test, p.value = 0.02) between the high-ethylene producer Etr1.1-NR1E and the low-ethylene producer ACO1 (**Fig. 2B**), due to increased root growth of poplar seedlings at the expense of photosynthetic tissue at high ethylene levels.

Overall, the metabolic profiles of root and shoot tissues from cuttings grown in sterilised soil showed no variation among genotypes, except for minor yet statistically significant alterations for a few metabolites in the roots of Etr1.1-NR1E genotype (Kruskall-Wallis, FDR correction, Dunn post-hoc, p.value ≤ 0.05) (**Fig. 2C**). Levels of amino acids (Glutamine, Asparagine, Threonine, Glutamate, Serine, Valine, Ornithine and Leucine), amounts of nitrogen compounds (Thymine), several sugars (arabitol and trehalose), as well as unknown compounds (14.05.292, 11.24.393, and 20.15.421) decreased specifically in the roots Etr1.1-NR1E genotype, while levels of stigmasterol increased (**Table S1-A**). Moreover, composition of root exudates after 30 days did not differ between genotypes (data not shown). In conclusion, the level of ethylene production led to changes in root-shoot ratio, but had no impact on root exudate composition or shoot metabolite profiles in the absence of microorganisms.

### Altered *in planta* ethylene levels deplete root metabolites in the presence of microbes and induce changes in shoot metabolites

Since ethylene signalling at the holobiont level is a regulatory cascade involving traits from both the plant and its associated microbiota (Ravanbakhsh et al. 2018), it was hypothesised that the plant development and metabolite profiles of root exudates, roots, and shoots in the presence of microbiota vary depending on the poplar genotype (low or high ethylene producer). Root exudates were collected from 4 to 30 days of growth to analyse the difference in composition between transgenic and WT-T89 poplars grown in microcosms with sterilised and natural soils taken from a poplar plantation. The root and aerial compartments after 30 days of growth were also sampled to determine their metabolomic composition and growth parameters (root-shoot ratio). The root-shoot ratio was similar within all the transgenic poplar lines, indicating that the presence of the microbiota mitigated the effect of ethylene overproduction on root-to-shoot ratio (**Fig. 3A**).

**Fig. 3.**
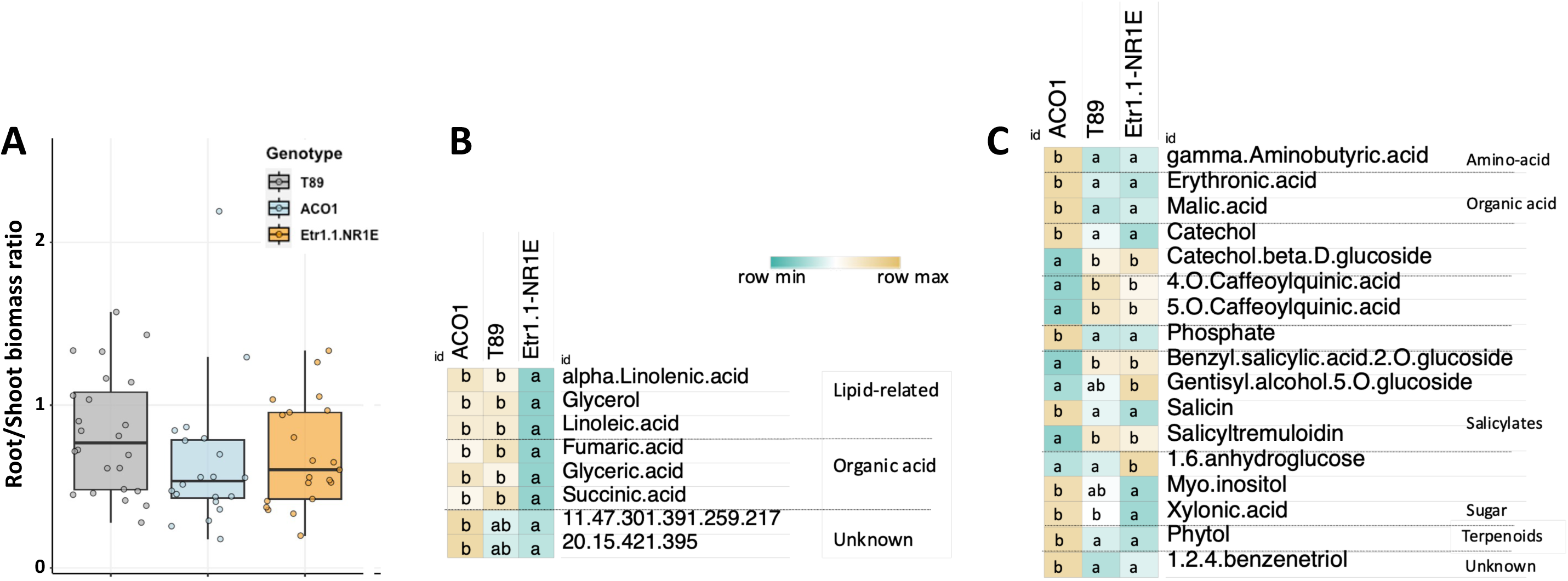
Impacts of the microbiota on the root-to-shoot ratio, and on the root and shoot metabolomes of poplar transgenic lines with modified ethylene regulation. (A) Root/shoot fresh biomass ratio of transgenic poplars and with wild type (LM, TuckeyHSD post-hoc test, p value > 0.05, n = 20-25) cultivated in natural soil. (B-C) Significant differences of metabolite abundances in (B) root and (C) shoot compartments between the transgenic line ACO1, Etr1.1.NR1E and wild type T89 in natural conditions after 30 days of growth (Kruskall-Wallis with FDR corrections, Dunn post-hoc test, p adjusted ≤ 0.05, n = 20-25, Table S1B and S1C). Each row represents the z-score of a metabolite in the three poplar genotypes. Unknown metabolites were named by retention time (min) followed by key mass-to-charge ratios (m/z) and putative metabolite classes are indicated

Root exudate compositions mainly differed between poplars cultivated in natural vs sterile soils, with a notable decrease in metabolite levels when microorganisms were present, as observed previously (Fracchia et al. 2024). Neither the Etr1.1-NR1E high-ethylene transgenic line, nor the low-ethylene transgenic line over-expressing ACO1 exhibited significant enrichment or depletion of any root exudates compounds at any time (Kruskal Wallis, p.value > 0.05). Differences of metabolites in both root and shoot were also mainly explained by the presence or absence of microorganisms (**Fig. 4**). However, the second principal component explained 20.1% of the total variance in roots and it was strongly correlated with the plant genotype, with the Etr1.1-NR1E genotype driving most of the changes (**Fig. 4A**). Comparison of metabolite levels between lines revealed three different patterns (**Table S1-B)**: metabolites with stable levels across all lines (89 compounds), metabolites showing the same trend of variation in both transgenic lines compared to WT (5 compounds) (**Fig. S1A**), and metabolites displaying opposite trends between the two transgenic lines (8 compounds) (**Fig. 3B**). We excluded from our analysis metabolites whose levels were significantly different from the WT in one line, but showed similar variation trends in both transgenic lines, considering that these variations were unlikely to be due to ethylene regulation. The same approach was similarly applied to microbial analyses. Considering this, eight metabolites were significantly depleted in the roots of Etr1.1-NR1E compared to ACO1 (Kruskal-Wallis test, FDR correction, Dunn posthoc test, p.value ≤ 0.05) (**Fig. 3B; Table S1-B**). Mainly reduced levels of lipid-related compounds such as alpha-linolenic acid, glycerol, and linoleic acid, along with decreased levels of intermediates of the tricarboxylic acid cycle, such as fumaric acid and succinic acid were observed. The microbiota also induced variation in shoot metabolites among the different genotypes (**Fig. 4B**), but in this case, ACO1 was the genotype explaining most of the variance. This was mainly driven by changes in salicylates and phenylpropanoids concentrations (**Fig. 3C; Table S1-C**). Seventeen shoot metabolites discriminated ACO1 and Etr1.1-NR1E genotypes, with opposite regulation. However, 26 shoot metabolites displayed the same tendency in reduction or increase in both genotypes (**Table S1-C, Fig. S1-B)**, suggesting that these changes are independent of ethylene level. In ACO1 aerial tissues, we observed a significant increase of GABA, phosphate, malic acid (**Fig. 3C; Table S1-C**) or other metabolites (galactonic acid, erythronic acid, sugars, and terpenoids). Finally, while salicin was significantly enriched in ACO1, other compounds related to *Populus* defence metabolites, including salicyltremuloidin, benzyl-salicylic acid-2-O-glucoside, precursor of phenylpropanoids significantly declined in comparison with ETR1.-NR1E and wild type plants (**Fig. 3C; Table S1-C**).

**Fig. 4.**
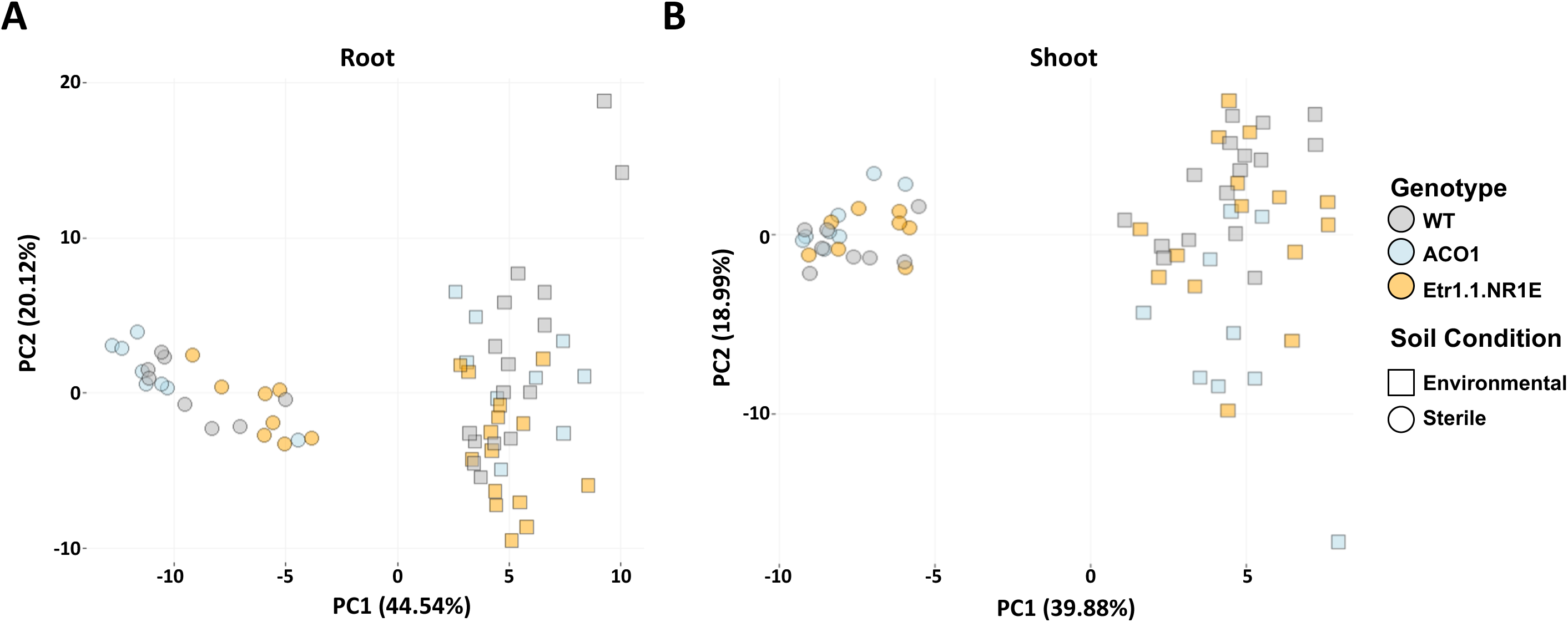
Principal component analysis of root and shoot metabolomes. Distribution of the metabolites detected in presence and absence of microorganisms in (A) root and (B) shoot poplar compartments after 30 days of growth. The two first principal components explain 64.66 and 58.87% of the observed variance in root and shoot compartments, respectively. Metabolites are mostly distributed regarding their abundance in natural and sterile soils rather than poplar genotypes (n = 3-5).

Altogether, our data suggest that ethylene overproduction (Etr1.1-NR1E line) in the presence of microorganisms negatively alters the composition of the root metabolomic profile, without impacting root exudates. The presence of microorganisms also suppresses the imbalance in root-shoot growth triggered by ethylene. On the other hand, low level of ethylene (ACO1) induces changes in shoot metabolomes (in presence of microorganisms) with an increase in basal metabolism (organic acids, sugars) and overall a decrease in defence compounds (salicylates and precursors of phenylpropanoids). Several hypotheses can explain the observed changes: (H1) these metabolite changes are the consequences of shifts in microbial community composition induced by ethylene level; (H2) these metabolite changes are the consequences from alteration in the activities of microbial communities; (H3) these metabolites changes are the results of the interaction between both shifts in microbial communities and activities.

### Bacterial and fungal communities are under the influence of poplar ethylene

To test these hypotheses, we analysed the microbial communities colonising the bulk soil, the rhizosphere, the roots and the shoots of WT-T89 poplar and transgenic lines altered in their ethylene production (Etr1.1-NR1E and ACO1) over time, from 4 to 30 days, combining a metabarcoding approach targeting bacterial (16S rRNA), Glomeromycetes (28S rRNA), fungal (Internal Transcribed Spacer: ITS) communities, and Confocal Laser Scanning Microscopy (CLSM) observations. CLSM analysis revealed six different types of fungal structures in the roots of poplar: ectomycorrhizae, arbuscular mycorrhizae (AM), single hyphae, coil-, glove-, and globular structures, as previously described (Fracchia et al. 2024; 2021) (**Fig. S2**). The relative abundance of coil structures was increased by 2-fold in ACO1 roots compared to WT-T89 and Etr1.1-NR1E lines (p.adj = 0.02, **Fig. 5A**). Furthermore, the proportion of hyphae detected outside the roots were 1.5x higher in the ACO1 lines than the WT (p.adj = 0.04, **Fig. 5B**).The metabarcoding approach allowed the detection, after rarefaction, a total of 1,675, 584 and 312 bacterial, fungal and glomerales Operational Taxonomic Units (OTUs), generated from 2,109,454, 1,107,547 and 1,020,453 high quality sequences, respectively. Time was the main driver structuring all of the microbial communities, regardless of the studied compartment, even though there was a significant effect of poplar transgenic lines on the structure of bacterial and fungal communities (**Table S2**). Saprotrophs dominated in all soil (35.17 ± 0.1%), rhizosphere (41.76 ± 0.14%), and root (30.86 ± 0.43%) compartments followed by EMF fungi (respectively 24.79 ± 0.19, 18.96 ± 0.3, and 28.08 ± 0.84%), where the saprotroph *Mortierella humilis* and the EMF *Inocybe curvipes* were the most abundant taxa (**Table S3; Table S4**). In shoots, saprotrophs also represented the major guild (37.57 ± 0.59%) with *Saitozyma podzolica* as the dominant taxa and endophytes such as *Clonostachys rosea* to a lesser extent (17.41 ± 1.09%) (**Table S3; Table S4**). Regarding bacterial communities, the unidentified *Candidatus udaeobacter* and *Acidobacteriae* subgroup2 dominated in the soil (respectively 12.94 ± 0.3 and 12.42 ± 0.18%) and the rhizosphere compartments (10.23 ± 0.34 and 10.12 ± 0.27%) (**Table S5**). The unidentified *Comamonadaceae* (11.97 ± 0.72%) and *Burkholderia Caballeronia Paraburkholderia* (8.49 ± 0.89%) represented the major bacterial taxa in roots, while the unidentified *Oxalobacteraceae* (25.71 ± 3.15%) and *Pseudomonas* (13.22 ± 2.13%) dominated the shoot compartments (**Table S5**). Finally, *Rhizophagus*, an unidentified OTU of Glomeromycotina and *Claroideo-Rhizophagus* were the dominant Glomerales genera in all soils (respectively 52.87 ± 2.19, 15.73 ± 1.44, and 13.3 ± 1.09%), rhizosphere (respectively 44.14 ± 2.51, 12.93 ± 1.69, and 12.51 ± 1.74%) and root compartments (respectively 60.81 ± 4.95, 10.88 ± 2.9, and 4.29 ± 1.48%) (**Table S6**).

**Fig. 5.**
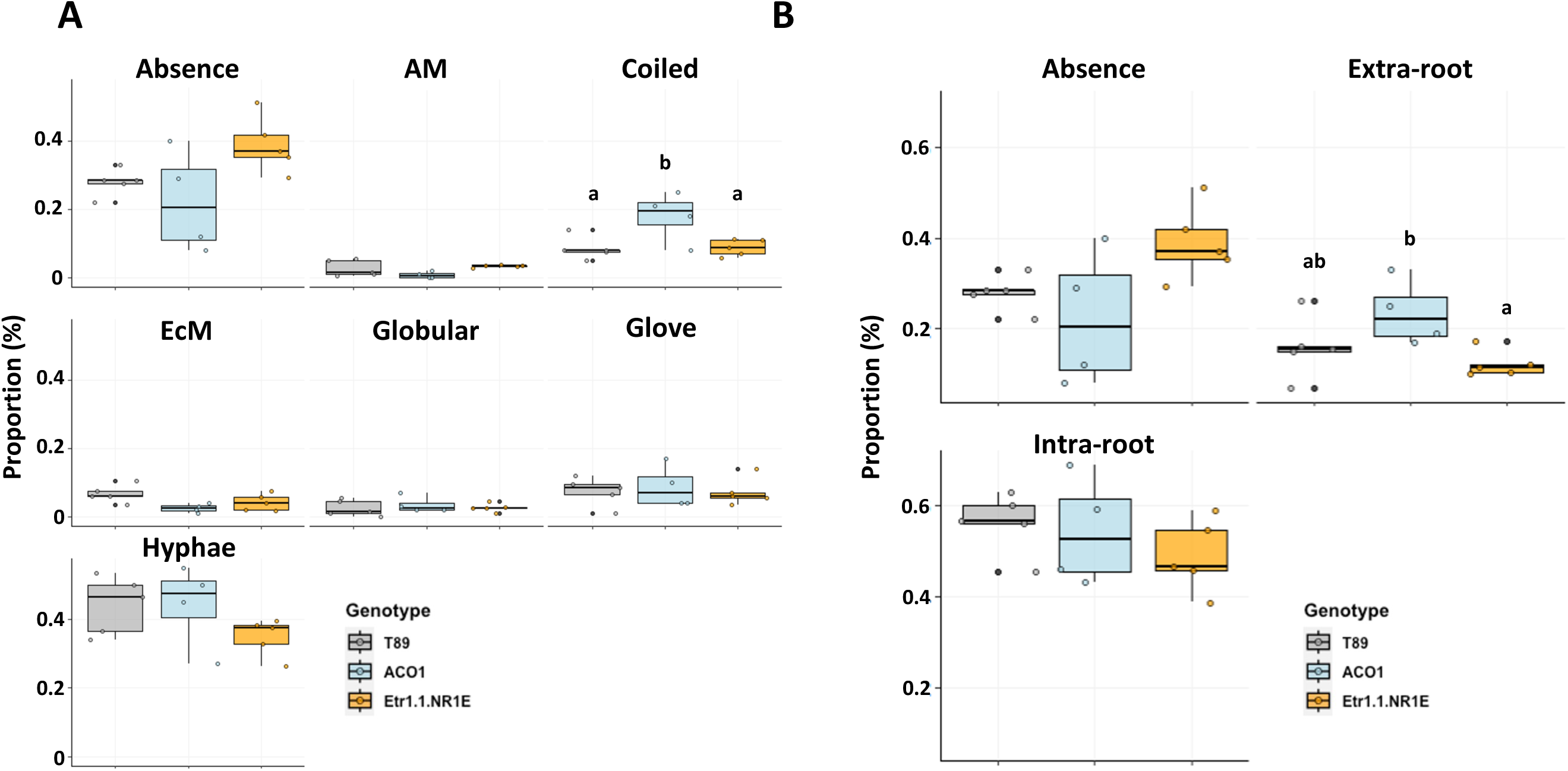
Relative abundances of fungal morphologies and their localisations in the root systems between transgenic and wild type poplars according to CLSM observations. (A) Relative abundance of the 6 different fungal structures detected in the roots of the 3 genotypes after 30 days of growth measured on 100 transversal transects per replicate. (B) Localisation of hyphae associated with the root systems after 30 days of growth. Letters indicate significant differences assessed by GLMM (beta distribution, Bonferroni correction, Tuckey HSD post-hoc test, p values ≤ 0.05, n = 4-5).

The overall structure of the community of Glomeromycetes fungi did not differ between poplar lines, regardless of the time or the sample compartment (soil, rhizosphere, roots) according to PERMANOVA analyses (data not shown). Only small differences were found between bacterial communities associated with the three genotypes: 5 and 8 % of the distance between communities were attributed to the genotypes in the rhizosphere and in the shoot associated bacterial communities (PERMANOVA analyses, p.value ≤ 0.05, **Table S2**). These community level changes were associated with a transient decrease of richness in the shoots of Etr1.1-NR1E line compared to the WT-T89 and ACO1 after 15 days (**Table S7**). Only 7% (17 out of the 239) bacterial genera detected showed a significant change in relative abundance between poplar genotypes (**Fig. S3; Table S5**). However, the relative abundance of only 8 bacterial genera detected in the soil, rhizosphere, and/or shoots significantly differed between the low and high-ethylene producers, and thus could be explained by ethylene levels (**Fig. S3; Table S5**). The bacterial genera *Ktedonobacteraceae HSB.OF53.F07* and *Aquisphaera*, as well as *Candidatus Koribacter* and *Granulicella*, were respectively enriched in the soil and rhizosphere of the high-ethylene producer Etr1.1-NR1E after 15 days of growth (**Fig. S3; Table S5**). Conversely, the genus *Acidipila-Silvibacterium* was enriched in the rhizosphere of the low ethylene producer ACO1 after 30 days of growth, whereas high ethylene levels favored the shoot colonisation of *Sphingomonas* by 7x, reaching 2.3% of the total read counts (**Fig. S3; Table S5**). Most of the changes were found in the fungal communities. Indeed, poplar genotype had a significant impact on fungal community structures as a function of time, in all sampled compartments, except roots (PERMANOVA analyses, p.value ≤ 0.05, **Table S2**). The strongest effect was found in the shoots for which genotype x time interaction explained up to 20% of the divergence between fungal communities, suggesting a time dependent influence of ethylene level on the fungal communities (**Table S2**). The alpha-diversity and the composition of the fungal communities differed between the three genotypes of poplar. The genotype ACO1, the ethylene-low producer, was characterised by a higher fungal richness in the roots after 30 days (**Fig. 6; Table S7**). Conversely, the Etr1.1-NR1E genotype, the ethylene-overproducer, showed a reduced fungal richness in the shoots compared to ACO1 after 30 days (**Fig. 6; Table S7**). At the level of trophic fungal guilds, a striking reduction of the relative abundance of saprotrophs in the shoots of Etr1.1-NR1E lines was observed in comparison with the WT and ACO1 lines after four days (Generalised Linear Model (GLM) with beta distribution, Bonferroni corrections, TukeyHSD post-hoc, p.value ≤ 0.05), shifting from 84% of the reads in average to less than 30% (**Fig. 7; Table S3**). However, this negative effect was lost over time, in conjunction with the overall decline in saprotrophs in shoots. To understand if this phenomenon was specific to some taxa or if it was more largely shared among members of trophic guilds, GLM was used to test if the relative abundance of the taxa varied in transgenic lines compared to WT-T89 lines at the three time points. Of the 255 fungal taxa identified in at least three replicates, 17% showed a positive or negative enrichment between poplar genotypes in at least one of the compartments investigated (**Fig. 8; Fig. S4; Table S4**). However, the level of ethylene could only explain the significant enrichment-depletion of 11 fungal species, and we speculate that the differences observed in the 31 fungal species remaining were either stochastic or due to the agro-transformation event (**Fig. 8; Fig. S4; Table S4**). Among these 11 fungal species, seven were affiliated to saprotrophs, two to endophytes, and only detected one EMF (**Fig. 8; Table S4**). During the early steps, low ethylene levels favored colonisation of three fungal taxa. The two low abundant saprotrophs, *Clavaria falcata* and *Dictyosporium toruloides*, were respectively enriched in the soil and rhizosphere of the low-ethylene producer ACO1, whereas the endophyte *Hyaloscypha finlandica* was enriched by 25x, reaching 2% of relative abundance in the shoot of ACO1 (**Fig. 8; Table S4**). This trend was reversed in the late colonisation stages, where high-ethylene levels exhibited a positive influence on colonisation of 8 fungal taxa in the soil, rhizosphere, and root compartments (**Fig. 8; Table S4**). One of the most striking results was the significant enrichment of the EMF *Tomentella ellisii* by almost 20x, reaching up to 3.3% of relative abundance in the soil of Etr1.1.NR1E after 30 days of growth (**Fig. 8; Table S4**). Similarly, the two saprotrophs *Mariannaea samuelsii* and *M. punicea* were enriched by more than 20x in the rhizoshere of the high-ethylene producer Etr1.1.NR1E, reaching 1% of relative abundance after 30 days (**Fig. 8; Table S4**). Finally, high-ethylene levels also advantaged the root colonisation of the saprotroph *Pezicula melanigena*, which was enriched by 4x in the roots of Etr1.1.NR1 (**Fig. 8; Table S4**).

**Fig. 6.**
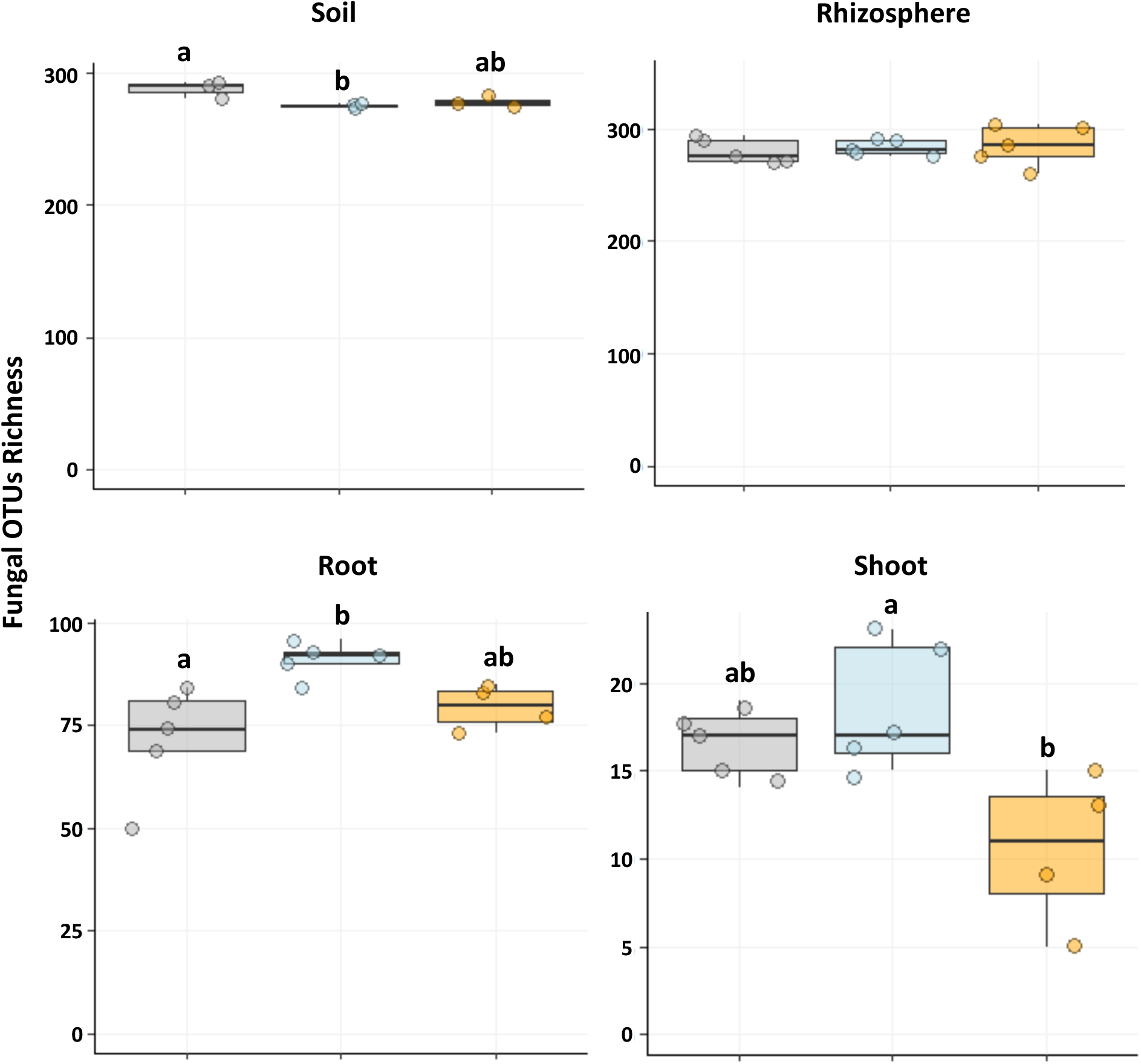
Richness of fungal OTUs associated with transgenic poplars after 30 days of growth. Richness of fungal OTUs detected in soil, rhizosphere, root and shoot of wild type and transgenic poplar after 30 days of growth. Letters indicate significant differences assessed by LM and corrected with Bonferroni, TuckeyHSD post-hoc test, p adjusted ≤ 0.05, n = 3-5.

**Fig. 7.**
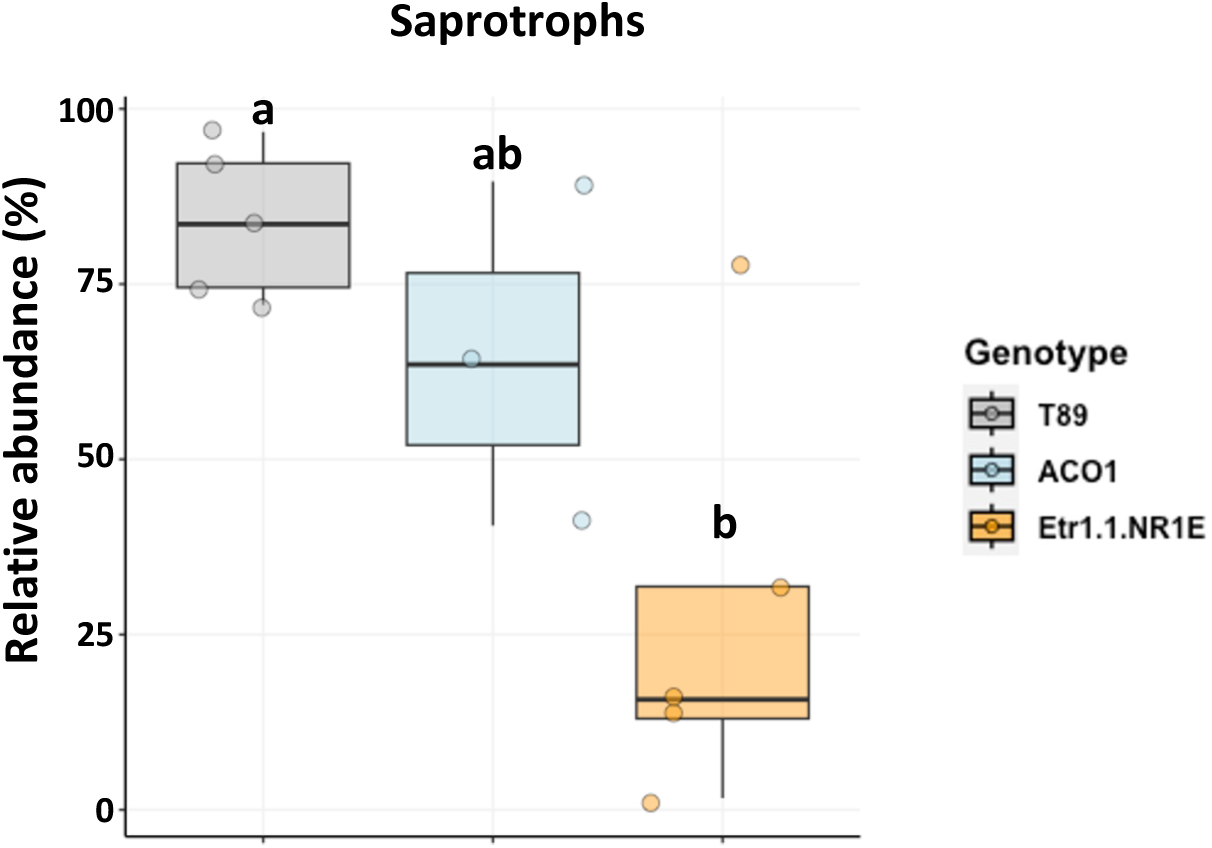
Relative abundance of saprophytes associated with transgenic poplars in shoots after four days of growth. Letters indicate significant differences assessed by GLM (beta distribution) and corrected with Bonferroni, TuckeyHSD PosHoc,, p adjusted < 0.05, n = 3-5.

**Fig. 8.**
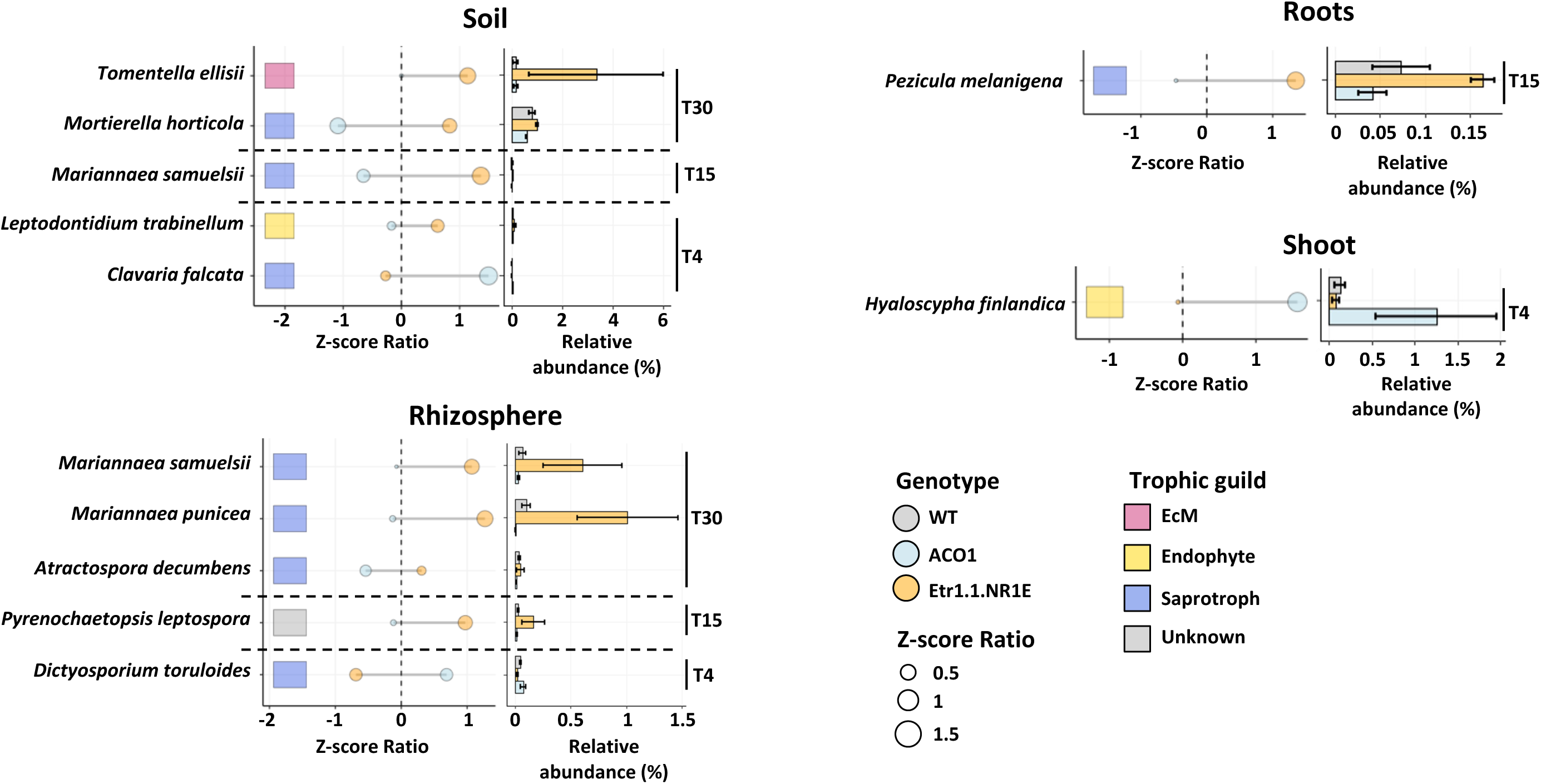
Ratio of the read counts and relative abundance of fungal species significantly different between low and high-ethylene producers in the soil, rhizosphere, root and shoot compartments over time. The left panel indicates the significant changes of fungal read counts, while the right panel indicate their relative abundance between low-ethylene, high-ethylene, and wild type poplar over 30 days of growth. Values in the left panel are given as the ratio between transgenic and wild type poplars of the reads counts expressed as z-score. Vertical dotted line represents the wild type as reference. Horizontal dotted line separates the three time points. Colors in the squares indicate the fungal trophic guilds. Significant differences were assessed by GLM with optimal family depending on the distribution of the read counts, FDR corrections, Tukey HSD posthoc test, p value ≤ 0.05, n = 3-5.

Taken together, these data suggest that ethylene levels mainly affect fungal rather than bacterial colonisation, and that the effects are a function of time and compartment. Low ethylene levels would favor the overall fungal richness, as well as the formation of coil structures in the roots and the density of external hyphae after 30 days, but elevated ethylene would reduce the early colonization of shoots by saprophytes. Nevertheless, species belonging to the same guild may exhibit disparate responses to the prevailing trend, suggesting a species-specific response to ethylene.

## DISCUSSION

When planted in soil, young sterile poplar seedlings are quickly colonised by microorganisms from the rhizosphere to the shoots, following a dynamic pattern in which early colonizers, dominated by saprotrophs, are slowly replaced by symbionts –AM, EMF and endophytes, and specific bacterial communities (Dove et al. 2021; Fracchia et al. 2024; 2021). This microbial colonisation is accompanied by massive changes in root exudation and in root and shoot metabolomes (Fracchia et al. 2024). In this study, we investigated the influence of the gaseous phytohormone ethylene on these dynamics. Ethylene is a central phytohormone which regulates the balance between growth and stress response, and elevated ethylene levels are generally a stress signal for plants, although the nature of the ethylene-mediated response is highly variable between plants (Ravanbakhsh et al. 2018). We show that, in poplar, the downregulation of ethylene production has a negligible impact on root exudation, and on the root and shoot metabolomes, when poplar seedlings are cultivated in sterile conditions (**Fig. S5**). However, when microorganisms are present, this downregulation gives rise to differential responses in roots and shoots, as well as an alteration of the microbial communities, particularly fungi.

### In poplar seedlings, AtEtr1-1 overexpression stimulated ethylene production compared to PttACO1 overexpression, in our experimental conditions

Quantification of ethylene production in young *in vitro* poplar seedlings revealed that the ethylene insensitive poplar lines Etr1.1-NR1E exhibited the highest level of ethylene, whereas the poplar lines over-producing PttACO1 showed no increase in ethylene production, but rather a tendency for its reduction. AtEtr1-1 encodes a mutated ethylene receptor (ETR1) that is active in both absence or presence of ethylene and activates the negative kinase regulator CTR1 (Azhar et al. 2023). In turn, the activation of CTR1 leads to ethylene signal inhibition. This latter would normally be alleviated by inactivation of the wild type ETR1 receptor when bound to ethylene (Camehl et al. 2010). The increased level of ethylene in ethylene-insensitive poplars is in accordance with previous literature. Ethylene-insensitive mutations (*ein1*, *ein2*, *eto1*, *eto3*) in *Arabidopsis* result in increased ethylene levels (Guzman and Ecker, 1990; Woeste, Ye, and Kieber 1999). While *ein2* and *ein3* mutants display high levels of ethylene in all vegetative tissues (Guzman and Ecker, 1990), the *eto1* and *eto3* mutants overproduce ethylene only in etiolated seedlings (Woeste, Ye, and Kieber 1999). In the former mutants, elevated levels of ethylene can likely be explained by the blocking of the inhibition feedback of ethylene biosynthesis. In the latter mutants, ethylene overproduction is due to a post-transcriptional action on the ACS (Woeste, Ye, and Kieber 1999).

However, according to the literature, the detection of less ethylene in the ACO1-overexpressor poplar plantlets was not expected. Love et al. (2009) first described those poplar lines and showed an increased level of ACO1 activity and ethylene production in the xylem and phloem of 2m-high poplar. In our case, we measured the ethylene produced by the whole *in vitro* grown poplar cuttings. The observed discrepancy might be linked to the developmental stage of the tree, the organ versus whole plant measurement, or the nutrient availability in the medium that can limit the level of the ethylene precursor ACC, the formation of which is the key regulatory step in ethylene biosynthesis. A few post-translational modifications of ACOs have been reported (Houben and Van de Poel, 2019; Xia et al., 2024) but studies of this type of regulation are still scarce. Using the same transgenic lines within an experimental framework more directly comparable to the current study, Plett et al. (2014) detected an increase of the expression of ERF genes in ACO1 lines, suggesting an overproduction of ethylene, while several ERF genes were down-regulated in Etr1.1-NR1E lines. These inconsistencies may rely on the presence of a dual regulation of some transcription factors ERF, independent of the level of ethylene via phosphorylation through the MPK3/6-cascade (Dubois, Van den Broeck, and Inzé 2018). Finally, Plett et al. (2014) quantified the expression of ERF genes when the transgenic lines were interacting with the EMF *Laccaria bicolor*. Yet, symbiotic interactions also play a role in the regulation of ethylene biosynthesis, and Splivallo et al. (2009) demonstrated the ability of EMF, such as the black truffle *Tuber melanopsorum*, to produce ethylene. Overall, the complex feedback loops and crosstalk between ethylene signalling and other pathways, could explain the differences of ethylene production observed between the two studies.

Ethylene affects both the growth and development of plants. Ethylene is most associated with the regulation of cell size, often restricting cell elongation, but it can also regulate cell division. For example, Love et al. (2009) showed a causal link between level of endogenous ethylene and the induction of cambium cell division and consequently, meristem growth, stimulating wood production in poplar. Exogenous ACC at low concentration stimulates lateral root primordia in many plant species, but it is also described as an inhibitor of primary root elongation, and lateral primordia initiation (while promoting the emergence of existing primordia) at high-concentration (Ivanchenko, Muday, and Dubrovsky 2008; Petricka, Winter, and Benfey 2012; Qin, He, and Huang 2019; Vanstraelen and Benková 2012). This resembles the hormesis effect: stimulation of a response at low doses and inhibition at higher doses (Agathokleous et al., 2019). At very low concentration, ethylene can stimulate leaf growth (Fiorani et al. 2002). We observed that root biomass was slightly higher in the ethylene-overproducer poplar transgenic line, while it decreased in ethylene-low producer line. Even though the concentration of ethylene measured in the present study is neither reaching an inhibition nor a stress threshold, our data indicate that high ethylene level induces root growth of poplar seedlings at the expense of photosynthetic tissues. The presence of the microbiota alleviated this modification with no change in the root-to-shoot ratio when poplars were cultivated in presence of microorganisms. These data are in accordance with one study demonstrating in *Arabidopsis* that bacterial microbiota can control root branching through the induction of the phytohormone ethylene response in the root and independently of nutrient availability and the auxin pathway (Gonin et al. 2023). This might be due to the ability of microorganisms themselves to produce and degrade ethylene, keeping an equilibrium of the root-shoot ratio between poplar genotypes. Indeed, several studies showed that the ability to produce ethylene is widespread among soil bacteria (Nagahama et al. 1992). Conversely, many rhizospheric fungi and bacteria possess ACC deaminase enzymes, which degrade ACC and ultimately lead to lower plant ethylene concentrations (Nascimento et al. 2014).

### Alteration of ethylene level does not impair root exudates but alters root metabolomes and the composition of the associated microorganisms

Under axenic conditions, we showed that ethylene overproduction slightly modulated the composition of root metabolites and particularly amino acids (**Fig. S5**). The latter, including glutamic acid -which is a hub in amino-acid metabolism and can be converted into glutamine and ornithine (and likely arginine that breaksdown to ornithine during TMS derivitization)- were depleted in the roots of the Etr1.1-NR1E line. The unique compound accumulating in roots of Etr1.1-NR1E poplars was stigmasterol, a sterol component of the plasma membrane and sterol-stress in plants (Valitova et al. 2024). These data may indicate that these transgenic ET-overproduced poplars have reduced amino acid metabolism and might be under stress. The root/shoot ratio of these transgenic lines indicate that roots developed more than the shoots, possibly in response to constitutive nutritional stress.

The most important metabolic changes between the ethylene-overproducer and ethylene- low producer were observed in roots and shoots in presence of microbiota (**Fig. S5**). The influence of root microbiota on changes in poplar root and shoot metabolites was previously reported (Fracchia et al. 2024; Kaling et al. 2018; Tschaplinski et al. 2014). Here, the overproduction of ethylene led to root depletion in lipids and organic acids involved in the TCA cycle, suggesting a potential reduction of root metabolic activity in the presence of microorganisms. Since microbial colonisation of roots was not modified in the Etr1.1-NR1E line according to CLSM and metabarcoding data (except for the enrichment of the saprotroph *Pezicula melanigena* after 15 days of growth), it could be speculated that elevated ethylene levels do not regulate the establishment of microbiota, but nutrient allocation to roots and microbial partners. Accordingly, ethylene signalling was found to be involved in the regulation of *Arabidopsis* root colonization by the growth promoting fungus *Pirifomospora indica* and to regulate the balance between beneficial and non-beneficial traits in the symbiosis (Camehl et al. 2010). Further analyses tracking nutrient fluxes between roots and their multiple partners would be necessary to validate this hypothesis. Nevertheless, our data suggest that elevated ethylene, at the level reached in our setup, does not inhibit root colonisation by symbiotic fungi (endophytic, arbuscular, and or ectomycorrhizal). Conversely, we observed an increased number of fungal species detected in the roots of the low-ethylene producing line ACO1, together with an enrichment in coil structures, without any change in the root metabolome, suggesting that low ethylene levels facilitate root fungal colonisation, particularly by endophytic coil-forming fungi, without increasing the metabolic burden on the host.

While ET-overproduction modified root metabolic profiles, it did not surprisingly impact root exudates, in both absence or presence of microorganisms. Numerous studies have highlighted the diversity of compounds in root exudates (e.g., McLaughlin et al. (2023), but also their plasticity and adaptive potential to shape the soil microbiome (Badri et al. 2013; Haichar et al. 2008; Hugoni et al. 2018; Micallef, Shiaris, and Colón-Carmona 2009; Zhalnina et al. 2018). Our data are not consistent with other studies on tomato indicating compositional changes in root exudates after the knockdown of an ethylene receptor and two Ethylene-responsive factors ERF-E2 or ERF-E3 encoding genes (Dyer et al. 2019; Fu et al. 2021). Root exudate alteration was associated with a modification of the rhizospheric community (Fu et al. 2021) and of roots attractiveness towards some species of parasitic nematodes, without impacting the interaction with the Plant Growth-Promoting Rhizobacterium *Bacillus subtilis* (Dyer et al. 2019). These discrepancies of responses between plant species highlight the fact that the role of root exudation in the regulation of root associated microbiota strongly differ between plant species as demonstrated for grass species (Guyonnet et al. 2018). Although the modulation of ethylene regulation did not alter the root exudation profile, the over-production of ethylene was associated with the enrichment in the rhizosphere and soil of four minor saprotrophic fungi, with two belonging to the *Mariannaea* genera, as well as the dominant EMF *Tomentella ellisii*, suggesting a direct effect of ethylene on specific fungal species via its diffusion in the rhizosphere. Conversely, only the chemoheterotroph bacteria *Acidiphila silvibacterium* was enriched in the rhizosphere with low ethylene level. The lack of change in poplar root exudation may explain the minor alteration of the bacterial community in the rhizosphere observed in our experiments, as we previously showed that bacterial communities exhibit a stronger response to root exudates than fungi (Fracchia et al. 2024).

### The alteration of ethylene level modulates shoot metabolomes along with microbial communities

Shoot metabolomes significantly differed between the three poplar lines, only in presence of microbiota. The ethylene-low producer line ACO1 displayed the strongest changes in shoot metabolites, with declines in defence compounds (such as salicylates and some phenylpropanoid conjugates, except salicin) and elevated levels of organic acid, sugars and phenolic compounds. On the other hand, the overproduction of ethylene (Etr1.1-NR1E) did not trigger strong changes in the shoot metabolites compared to WT. As for the roots, these metabolic changes did not reflect microbial colonisation patterns, which showed the opposite trend (**Fig. S5**). The shoot microbiota of the Etr1.1-NR1E line were colonized by a lower number of fungal species, in particular less saprotrophic fungi. These results may reflect a dual effect of ethylene, which at high concentrations inhibits the development of some fungi, while at low concentrations regulates the activities of the microbiota without affecting their ability to colonise tissues. Ethylene has been shown previously to directly regulate the activity of the mold *Aspergillus parasiticus* without affecting the growth of the fungus (Gunterus et al. 2007), while other studies have shown differential inhibitory effects of ethylene depending on ethylene level and fungal species (Pristijono et al. 2018). This dual effect may not be that surprising knowing that ethylene has both stimulatory and inhibitory effects on plant growth depending on tissue and concentration (Dugardeyn and Van Der Straeten 2008). Little data are available regarding the role of ethylene on the regulation of non-pathogenic leaf microbiota. Bodenhausen et al. (2014) found that *Arabidopsis ein2* mutants altered in ethylene signalling were more colonised by the *Variovorax* bacteria in a synthetic community experiment assembling 7 bacterial species, but to the best of our knowledge, no data are available regarding fungal communities. Our results suggest that fungal communities may be more responsive than bacterial communities, and thus merit more attention.

## CONCLUSION

The up- or downregulation of ethylene production and signalling in transgenic poplar partially impaired the establishment of fungi and to a lesser extent bacteria, in the rhizosphere-root-shoot continuum. The effects were accompanied by contrasting poplar responses at the metabolomic level. While ethylene did not influence root exudation, regardless of its direct diffusible influence on microbial communities, the root metabolome reflected a reduced metabolic activity despite colonisation by symbiotic fungi. This suggests a potential role of ethylene in regulating nutrient fluxes towards symbionts. Furthermore, our data indicate a possible dual role for ethylene, where its levels may either promote or inhibit microbial growth and activity, depending on the concentration, as in hormesis effects. Knowing that strong interconnections exist with jasmonic acid and salicylic acid pathways, future research should investigate the potential role of these phytohormones in the control of poplar colonisation by commensal microorganisms and of their activities.

## MATERIALS AND METHODS

### Biological material

All plant material was derived from hybrid aspen *Populus tremula* x *tremuloides* Michx., clone T89. Three different lines produced by Love et al. (2009) were used in this study, including a 35S::PttACO1 line over-expressing the ACO1 gene involved in the last step of ethylene biosynthesis, and two independent lines of 35S::At*etr1.1*, that overexpress an *etr1-1* mutant allele of *A. thaliana* leading to a reduced perception of ethylene. *Populus tremula* x *tremuloides* T89 (wild-type WT) was used as control for all experiments. The over-expression of the transgenes in root and aerial compartment was verified for each transgenic line by real-time quantitative polymerase chain reaction (qRT-PCR). All poplar lines were propagated aseptically on Musharige-Skood medium (MS) and were cultivated at 24°C in a growth chamber (photoperiod 16h, light intensity 150 μmol.m^-2^.s^-1^)

Plantlets were produced by planting internodes with a single bud on MS supplemented with indole-3-butyric acid (IBA) (2 mg.L^-1^) during 1 week before being transferred on MS for 2 weeks until root and shoot development. This development growth procedure was used to generate rooted plantlets for all experiments.

### Quantification of ethylene production

After *in vitro* poplar development growth as described above, a total of 60 plants per transgenic line and control were grown on MS between two sterile pretreated cellophane membranes (Felten et al. 2009). Cellophane membranes were boiled in EDTA (1.g.L-1), rinsed with deionised water twice, and then autoclaved twice in water. After 3 weeks, we harvested the poplar seedlings and transferred 3 plants per line into 40 ml vials containing 5 ml of sterile water, giving a total of 20 vials per line considered as 20 replicates. Vials were hermetically sealed with a polytetrafluoroethylene (PTFE) silicone septum, optimised for organic volatile compounds (VOC) analyses (Thermo Scientific™). After 6 hours of confinement, 1 ml of gas was collected with a 1 ml syringe mounted with a 0.45 x 13 mm needle and transferred into 2 ml HPLC vials (vials Interchrom 282662, stoppers Interchrom CH227140) after removing 1 ml of gas from the HPLC vials. Vials were directly shipped and immediately injected in a gas chromatograph (GC) for analyses after receipt, as described previously (Trapet et al., 2016). Briefly, one mL of headspace gas was sampled and analysed by GC (Agilent 7820A) using a 2 m x 3 mm 80/100 alumina column, an injector at 110°C, N2 vector gas in an isocratic oven at 70°C, and a flame ionization detector at 250°C. The calibration was run using ethylene gas standards with a similar sampling schedule, as described above, to take the dilution into account.

### Soil collection and preparation

To reconstruct a forest-like microbial inoculum, soil (clayey loamy soil type) was collected from an 18-year-old poplar stand planted with *Populus trichocarpa* × *deltoides* and located in Champenoux, France (48 4592499N/6 2192499E) after pruning of brambles and adventitious plants and litter removal with a rake, as previously described (Fracchia et al. 2024). The first soil horizon (0 to 20 cm) was collected over an area of 1 m^2^ under 5 different trees. The soil was dried at room temperature for a week, and then sifted at 2 mm before being transferred to microcosms (see below). Three subsamples of 20 g each were harvested and stored at - 80°C until further analyses. To decipher the effects of ethylene production alteration on poplar root exudates and metabolomes, a subset of 50 kg of soil distributed in sealed bags of 200g each was sterilised by gamma radiation (45-65 kGy, Ionisos, France). Gamma irradiated soils were conserved after irradiation at room temperature for 3 months to allow for inactivation before use. Physico-chemical properties of soil are available in Fracchia et al., (2024).

### Plant growth and sampling procedure

To investigate the role of ethylene in structuring microbial communities, 200 g of soil were dispersed into 1500 cm^3^ boxes closed with filtered lids (OS 140 Box, Duchefa-biochimie), and soil was maintained at 75% humidity. We transferred 2 homogenous *in vitro* seedlings (1 cm long for aerial parts and 1-2 cm long root systems) from the 4 poplar lines in pots containing the environmental soil mentioned above. Each pot was closed with a filtered cover allowing gas exchange and we wrapped the bottom (approximately 1/3 of the pot) with aluminium foil to prevent algal and moss development. We cultivated the plants at 24 °C in a growth chamber under the same conditions mentioned above (photoperiod 16h, light intensity 150 μmol.m^-2^.s^-1^). In total, we grew over 4, 15 and 30 days, 300 plants (100 plants / 50 pots per line) distributed among 150 pots. Regarding microbial community analyses, at each time point, and for the 3 lines, we collected bulk soil, rhizosphere, root and shoot compartments from 5 plants (**Fig. 1**). We separated aerial and root parts, and collected the rhizosphere by pouring the root systems with adherent soil in 15 ml falcon tube containing 2 ml sterile 1X phosphate-buffered saline (PBS; 0.13 M NaCl, 7mM Na2HPO4, 3mM NaH2PO4 [pH 7.2]). After removing the root systems, we briefly vortexed the falcon tubes containing the rhizosphere and centrifuged them for 10 minutes at 4000 rpm. Then, we removed the supernatant to only keep rhizosphere samples. Finally, we washed the root compartments in sterile water to remove remaining soil particles. Soil, rhizosphere, aerial and root samples were frozen in liquid nitrogen and stored at -80 °C until DNA extraction. We also harvested *in vitro* poplars to confirm their axenic status prior to planting (time point T0).

### Collection of root exudates

To characterize root exudates, we collected 15 plants at each time point for all poplar lines from both gamma-irradiated or non-irradiated soils as described in Fracchia et al. (2024) (**Fig. 1**). We washed the roots in sterile water to remove adherent soil particles. Three poplar seedlings were transferred in 20 ml vials where roots were immersed in 5 ml of hydroponic solution, giving a total of 5 vials per genotype considered as 5 replicates. We let the root systems exude for 4h at 24 °C before collecting poplar seedlings. We collected the root exudates in hydroponic solution after filtering using HPLC compatible filters (Acrodisc® 25mm syringe filter with 0.2µm WWPTFE membrane) and stored them at -80 °C before further analyses. Five plants used for root exudates analyses were used to further characterize fungal colonisation by CLSM imaging. To do so, we separated aerial and root parts, and fixed the root compartment in Eppendorf tubes containing 1 ml of 3% paraformaldehyde (PFA) at 4 °C overnight for further staining and CLSM observations.

### Metabolite analyses

To detect if the alteration of ethylene regulation in *Populus* modulates the metabolomic profile of aerial and root tissues, we analysed the metabolite composition for both shoot and root compartments after 30 days of growth (**Fig. 1**). In addition, we followed the exudate composition from 4 to 30 days of growth between the 4 poplar lines. Concerning root exudates, we purified them using Sep-Pak C18 cartridges (Waters^TM^) in order to remove the salts contained in the hydroponic solution. In brief, we conditioned the column by loading 700 μl (one volume) 7 times with 100% acetonitrile. We then equilibrated the column with 7 volumes of H_2_O before loading 2 ml of exudates. We washed the columns with 5 volumes of water and eluted our samples in three steps with an acetonitrile gradient ranging from 20%, 50% and 100%. Finally, we weighed the lyophilised root exudates, and analysed their content by GC-MS. For root and shoot metabolomic, we harvested after 30 days of growth between 20 to 25 seedlings from the 4 lines. We weighed root and aerial part before lyophilisation and weighed them again. We then pooled aerial and root parts corresponding to each poplar lines to obtain between 9 to 15 replicates of dry material ranging between 25 to 100 mg for both compartments. We ground the dry shoot and root material using metal beads and a tissue-lyzer before analysing their metabolomic composition GC-MS.

Untargeted metabolite levels were determined from lyophilized roots and aerial compartments as described in (Tschaplinski et al. 2012). To ensure complete extraction, powdered and freeze-dried material (∼25 mg) was twice extracted overnight with 2.5 mL of 80% ethanol, sorbitol (75 µL (L) or 50 µL (R and Myc) of a 1mg/mL aqueous solution) was added to the first extract as an internal standard to correct for subsequent differences in derivatization efficiency and changes in sample volume during heating. The extracts were combined, and 500-µL (L) or 2-mL (R and Myc) aliquots were dried under nitrogen. Metabolites were silylated to produce trimethylsilyl derivatives by adding 500 µL of silylation-grade acetonitrile to the dried extracts followed by 500 µL of N-methyl-N-trimethylsilyltrifluoroacetamide with 1% trimethylchlorosilane. After 2 days, a 1-µL aliquot was injected into an Agilent Technologies (Santa Clara, CA) 7890A/5975C inert XL gas chromatograph / mass spectrometer (MS) configured as previously described (Tschaplinski et al. 2012). The MS was operated in electron impact (70 eV) ionization mode using a scan range of 50-650 Da. Metabolite peaks were quantified by area integration by extracting a characteristic mass-to-charge (m/z) fragment with peaks scaled back to the total ion chromatogram using predetermined scaling factors and normalized to the extracted mass, the recovered internal standard, the analysed volume and the injection volume. The peaks were identified using a large in-house user-defined database of ∼2600 metabolite signatures of trimethylsilyl-derivatized metabolites and the Wiley Registry 10th Edition combined with NIST 2014 mass spectral database. The latter commercial database has recently been upgraded to the Wiley 12^th^ Edition combined with NIST 2020 and all unknowns were run through these updated databases. The combination of these databases allowed us to accurately identify a much larger fraction of the observed metabolites compared to most analytical labs.

### Microbial community analyses

#### DNA extractions and amplifications

For soil and rhizosphere, we extracted DNA from 250 mg of material using the DNeasy PowerSoil kit according to the protocol provided by the manufacturer (Qiagen). For root and aerial compartments, we used 50 mg of ground plant material (less than 50 mg for certain root systems before transplanting and after four days of growth) to extract DNA using DNeasy Plant Mini kit following the manufacturer protocol (Qiagen). We quantified DNA concentration using a NanoDrop 1000 spectrophotometer (NanoDrop Products) and we normalised DNA extraction to the final concentration of 10 ng.µl^-1^ for soil and rhizosphere samples and 5 ng.µl^-1^ for root and shoot samples.

To maximize the coverage of bacterial 16S rDNA and fungal ITS2 rDNA regions, we used a mix of forward and reverse primers as described in (Mangeot-Peter et al. 2020). Regarding bacterial communities, we used a combination of 4 forward and 2 reverse primers in equal concentration (Fracchia et al. 2024), targeting the V4 region of the 16S rDNA. For fungal communities, we used 6 forward primers and one reverse primer in equal concentration targeting the ITS2 rDNA region (Fracchia et al. 2024). To avoid the amplification of plant material, we used a mixture of peptide nucleic acid (PNA) probes inhibiting the plant mitochondrial (mPNA) and chloroplast DNA (pPNA) for 16S libraries, and a third mix of PNA blocking the plant ITS rDNA (itsPNA) (Lundberg et al., 2013).

Knowing that our amplification protocol for fungal library is not efficient to detect Glomeromycota, we adapted a protocol from Brígido et al. (2017) to target specifically the 18S DNA region of this ecological guild. In brief, we applied a two steps PCR, where we used the LR1 (10 µM) and NDL22 (10 µM) primers (Van Tuinen et al. 1998), which amplify the large ribosomal subunit (LSU) DNA for the first PCR amplification. In the second PCR, we used the FRL3 (10 µM) and FRL4 (10 µM) primers (Gollotte, Van Tuinen, and Atkinson 2004) that specifically amplify the LSU-D2 rDNA of AM fungi. PCR conditions and programs were done according to Fracchia et al. (2024).

For the three libraries (16S, ITS, 18S) we used PCR mix without addition of DNA (negative control) and known fungal or bacterial communities (mock communities) as quality controls. The amplicons were visualised by electrophoresis through a 1% agarose gel in 1X TBE buffer. We purified PCR products using the Agencourt AMPure XP PCR purification kit (Beckman Coulter) following the manufacturer protocol. After DNA purification, we quantified PCR products by Qubit^®^2.0 fluorometer (Invitrogen) and performed new PCRs for samples with a concentrations lower than 2,5 ng.µl^-1^. Samples with DNA concentrations higher than 2,5 ng.µl^-1^ were sent for tagging by PCR and MiSeq Illumina next-generation sequencing (GenoScreen).

### Microbial Sequence processing

After quality controls, sequences demultiplexing and barcodes removal, we processed fungal, bacterial and glomerales sequences using FROGS (Find Rapidly OTU with Galaxy Solution) (Escudié et al. 2018) implemented on the Galaxy analysis platform (Afgan et al. 2016). Sequences were clustered into OTUs based on the iterative Swarm algorithm, and then chimeras and phiX contaminants were removed. As suggested by Escudié and collaborators (2018), we removed OTUs with a number of reads lower than 5.10^-5^ percent of total abundance. We then discarded fungal sequences not assigned to the ITS region using the ITSx filter implemented in FROGS and we affiliated fungal sequences using the UNITE Fungal Database v.8.3 (Nilsson et al. 2019), and bacterial sequences using SILVA v.138.1 database. We considered OTUs with a BLAST identity lower than 90% and BLAST coverage lower than 95% as potential chimeras and removed these sequences from the dataset. As well, we removed sequences affiliated with chloroplast and mitochondria. As a means to optimise the analyses of microbial community structures and diversity, we applied different rarefaction thresholds depending on compartments and communities. Regarding microbial structure analyses, we rarefied the number of sequences depending on the studied compartment but without considering time to be able to account for the time effect in shaping microbial structure (**Table S8**). Thus, microbial structures associated to the distinct compartments were analysed separately. On the other hand, for alpha diversity and microbial composition analyses, we rarefied microbial sequences for each time and compartment (**Table S8**). As a result, we analysed the alpha diversity and the composition of microbial communities for each time and compartment. We combined two databases, FUNGuild (Nguyen et al. 2016) and FungalTraits (Põlme et al. 2020) to classify each fungal taxa into an ecological trophic guild. We applied a confidence threshold only keeping “highly probable” and “probable” affiliated trophic guilds and we assigned the other fungal taxa as “Unknown”.

### Confocal laser scanning microscopy observation

To investigate if transgenic poplar altered in ethylene regulation showed a modulation of fungal spatio-temporal root colonization, we performed CLSM observations after staining fungal structures and poplar root systems. We adapted the staining procedures of root systems from (Vierheilig, Schweiger, and Brundrett 2005). In brief, we washed the fixed root systems three times in one volume 1X PBS. We cleared the root systems collected after 15 and 30 days of growth in 20% KOH during 2h at 90°C. Then, we washed the root systems 3 times in 1X PBS and we incubated them overnight in the dark in 1 ml of 1X PBS containing 10 µl of 10 mg.ml^-1^ WGA-Alexa Fluor 488 (Thermo Fisher Scientific, Waltham, MA), a specific marker of the chitin of fungal cell walls. After fungal staining, we washed the root systems 3 times in 1X PBS and incubated them in 1 ml 1X PBS supplemented with 10 µl of 10 mg.ml^-1^ propidium iodide (PI), a DNA intercalating agent that also stains plant cell walls. After 30 min, we washed the root systems 3 times in 1X PBS before mounting them between slide and cover slip with a few drops of 20% glycerol. All root samples were observed with a Zeiss LSM 780 confocal laser scanning microscope (Zeiss International) using 10X, 20X, and 40X objectives. WGA-Alexa Fluor 488 was excited using a 488-nm excitation wavelength and detected at 500 to 540 nm, whereas a 561-nm excitation wavelength and detection at 580 to 660 nm were used for propidium iodide. Maximum intensity projections were performed using ZEN software with a z-stack width of 30 to 50 μm. We quantified fungal root colonization by adapting the grid-intersect method developed by (Fracchia et al. 2023; McGONIGLE et al. 1990).

### Statistical analyses

We used the R software v.4.3.0 (R Core Team. 2023) and RStudio v.2023.03.1 (RStudio Team, 2023) to compute all data analyses, statistics and graphical representations, and we generated all figures using the package ggplot2 v.3.4.4 (**Wickham et al., 2016**). Unless mentioned otherwise, we considered p.values ≤ 0.05 as significant. We used the vegan package v.2.6.4 to analyse the beta diversity of the microbial communities (**Oksanen et al., 2022**). We performed linear models (LM), linear mixed effect models (LMM) and generalised linear models (GLM) using respectively the *lm*, *lmer,* and *glm* functions from the lme4 package v.1.1.35.1. The generalised mixed effects models (GLMM) were performed using the *gamm* function from the mgcv package v.1.9.0. For microbial composition, we used the *gamlss* function from the gamlss package v.5.4.22 which allows for zero-inflated models, and the *AIC* function to calculate the Akaike information criterion (AIC). After checking the normal distribution of the data, we assessed the significant differences of ethylene production and root/shoot ratio between poplar transgenic lines and wild type with linear models followed by TukeyHSD post-hoc tests. We assessed the differences in metabolomic profiles between transgenic lines and wild type using Kruskal-Wallis tests, corrected by FDR, followed by a Dunn post-hoc test. Differences in fungal community structures between genotypes were tested using permutational multivariate analysis of variance (PERMANOVA) based on Bray-Curtis dissimilarity matrix distances using the *adonis2* function from the vegan package. We assessed the difference of microbial richness between genotypes at the different sampling times using LM, followed by TukeyHSD post-hoc test. We investigated the variations of the relative abundance of fungal trophic guilds between genotypes using GLM with a beta distribution and FDR corrections, followed by TukeyHSD post-hoc test. We tested the difference of microbial abundance between genotypes using the number of reads after rarefaction at each time and compartment and at different taxonomy levels, from the OTU to the family. We assessed the reads distribution of each taxon at a given time and compartment, and applied GLM with four distinct laws used for counts: negative binomial (NBI), zero-inflated negative binomial (ZINBI), Poisson (PO) and zero-inflated poisson (ZIP). We tested the quality of all four models by comparing the two probability distributions by plotting the quantile-to-quantile relationships (QQ plot). We then calculated the AIC, and retained the model with the lowest AIC value to investigate the influence of poplar ethylene production on microbial taxa. Finally, we assessed the different colonisation between genotypes observed by CLSM using GLMM with a beta distribution.

## Declarations

**Ethics approval and consent to participate** noy-applicable.

**Consent for publication** not-applicable.

### Availability of data and materials

Raw data were deposited in the NCBI Sequence Read Archive (SRA) under SRA accession numbers SRR26346063 to SRR26346115, and SRR32461720 for the 16S data, SRR26286627 to SRR26286662, and SAMN46983216 to SAMN46983270 for ITS data and SRR26286064 to SRR26286106 and SRR32461851 to SRR32462781 for 28S data (project PRJNA1017804).

### Competing interests

The authors declare that they have no competing interests.

### Funding

FF was supported by “Contrat Doctoral” from the Lorraine Université d’Excellence. This research was sponsored by the Plant-Microbe Interfaces Scientific Focus Area in the Genomic Science Program, the Office of Biological and Environmental Research in the U.S. Department of Energy Office of Science. Oak Ridge National Laboratory is managed by UT-Battelle, LLC, for the U.S. Department of Energy Office of Science (DE-AC05-00OR22725), LABEX ARBRE (ANR-11-LABX-0002-01).

### Authors’ contributions

FF, CVF and AD designed and coordinated the research and the experimental design. Poplars *in vitro* cultures were produced and maintained by FF and FG. Sampling was conducted by FF, SW and FG. CLSM observations were performed by FF and MB. DNA extractions and PCR amplifications were done by FF. CC quantified the ethylene production and TT and NE conducted the metabolomic analyses. Data analysis was performed by FF, CVF, and AD. FF, AD, and CVF wrote and FF, AD, CVF, CC, and TT revised the manuscript. All authors approved the final version of the manuscript

## Acknowledgements

Not applicable

## Notes

### Competing Interest Statement

The authors have declared no competing interest.

