## Supplementary figures and images for "Manipulation of *in planta* ethylene levels modulates the metabolome of *Populus tremula x tremuloides* in a microbial-dependent manner"

### Supplemental Figures

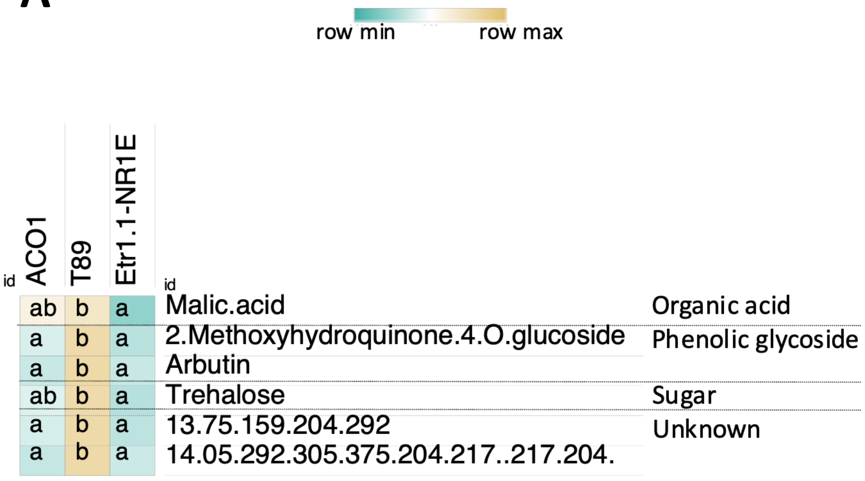

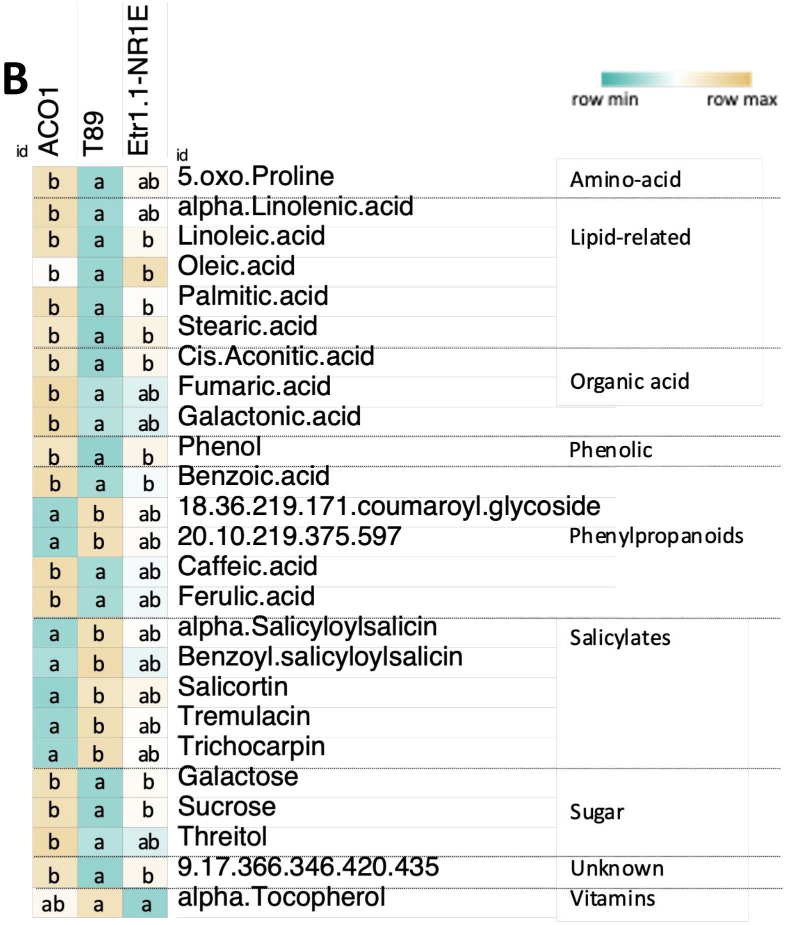

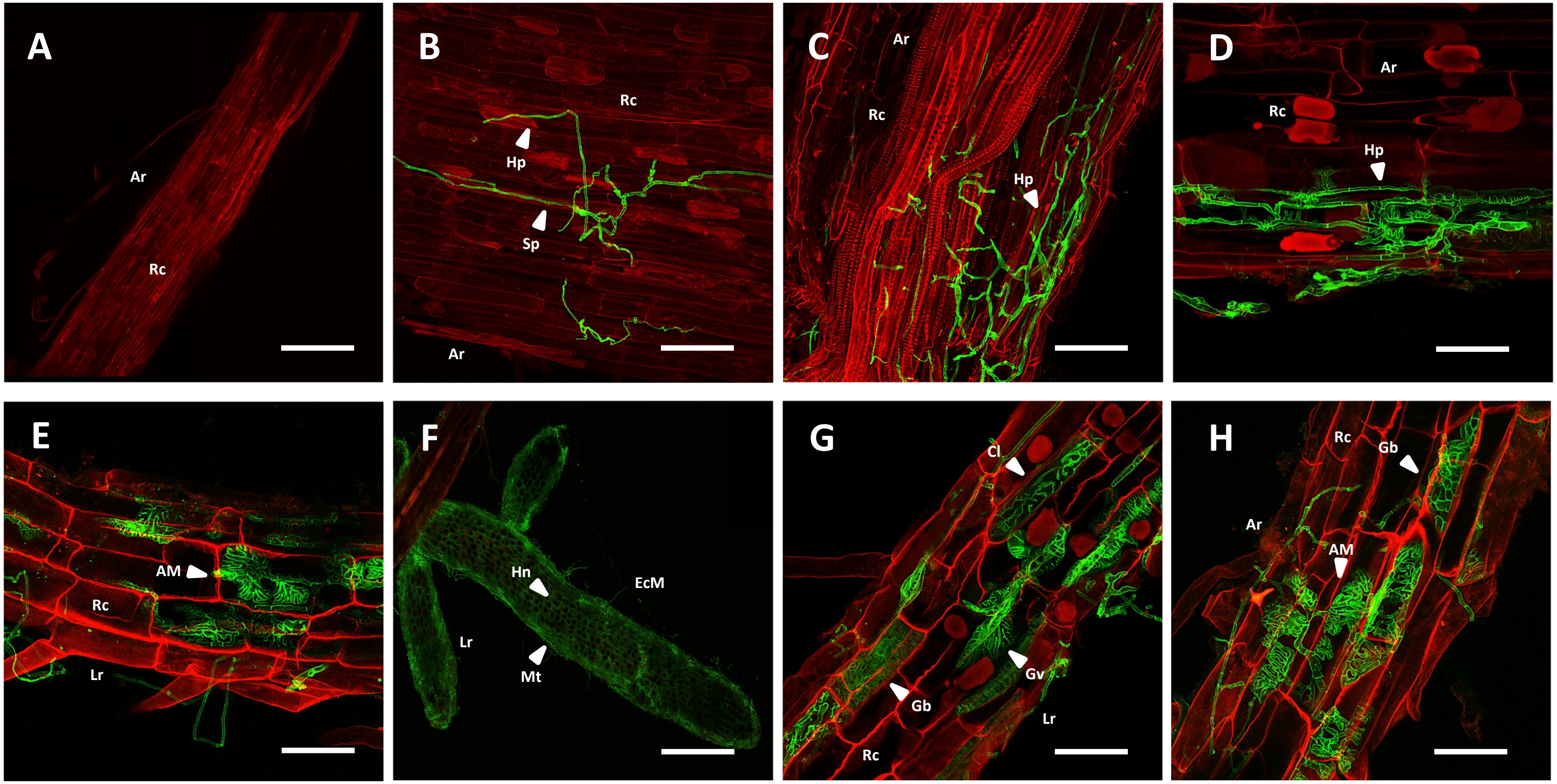

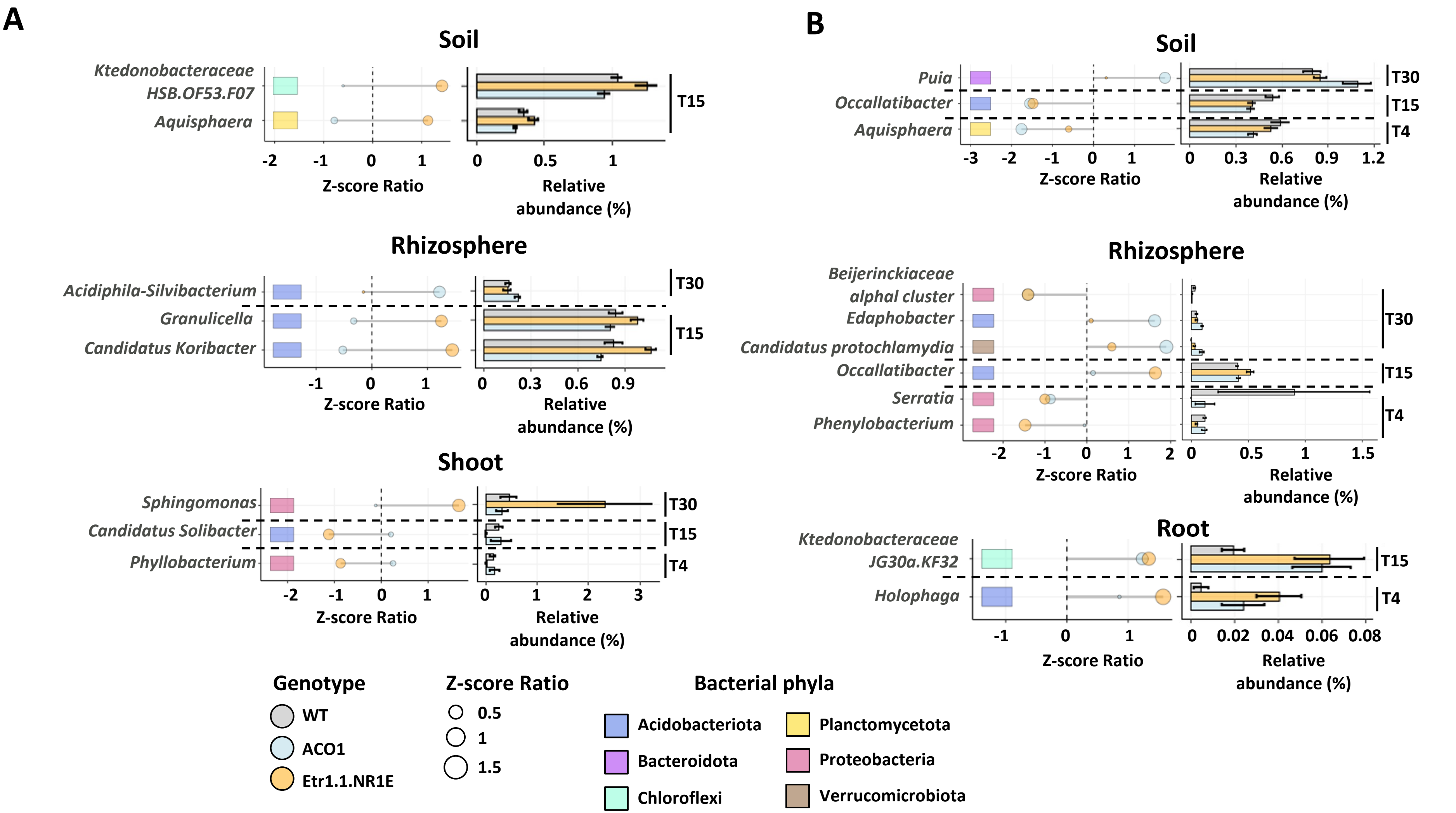

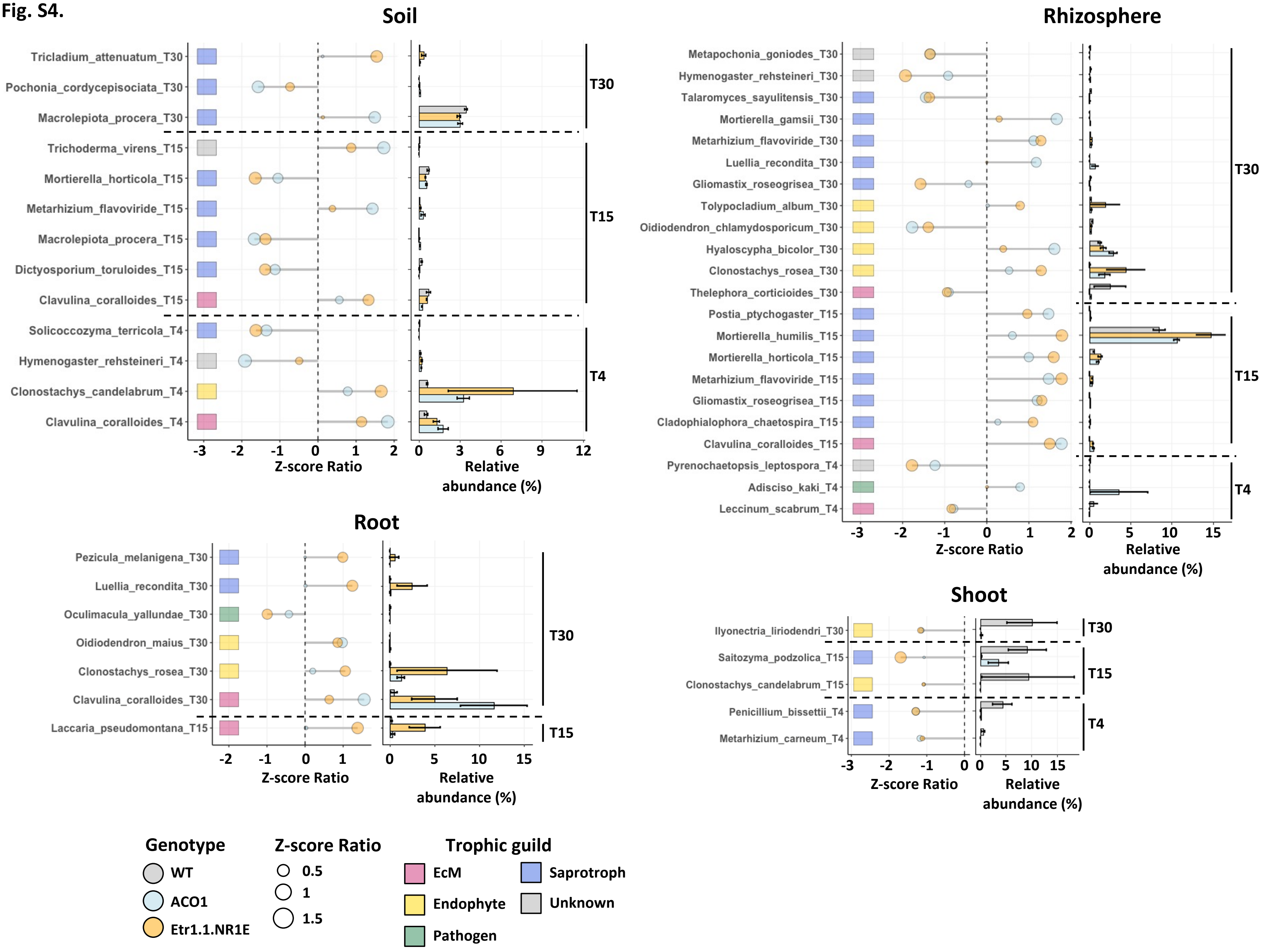

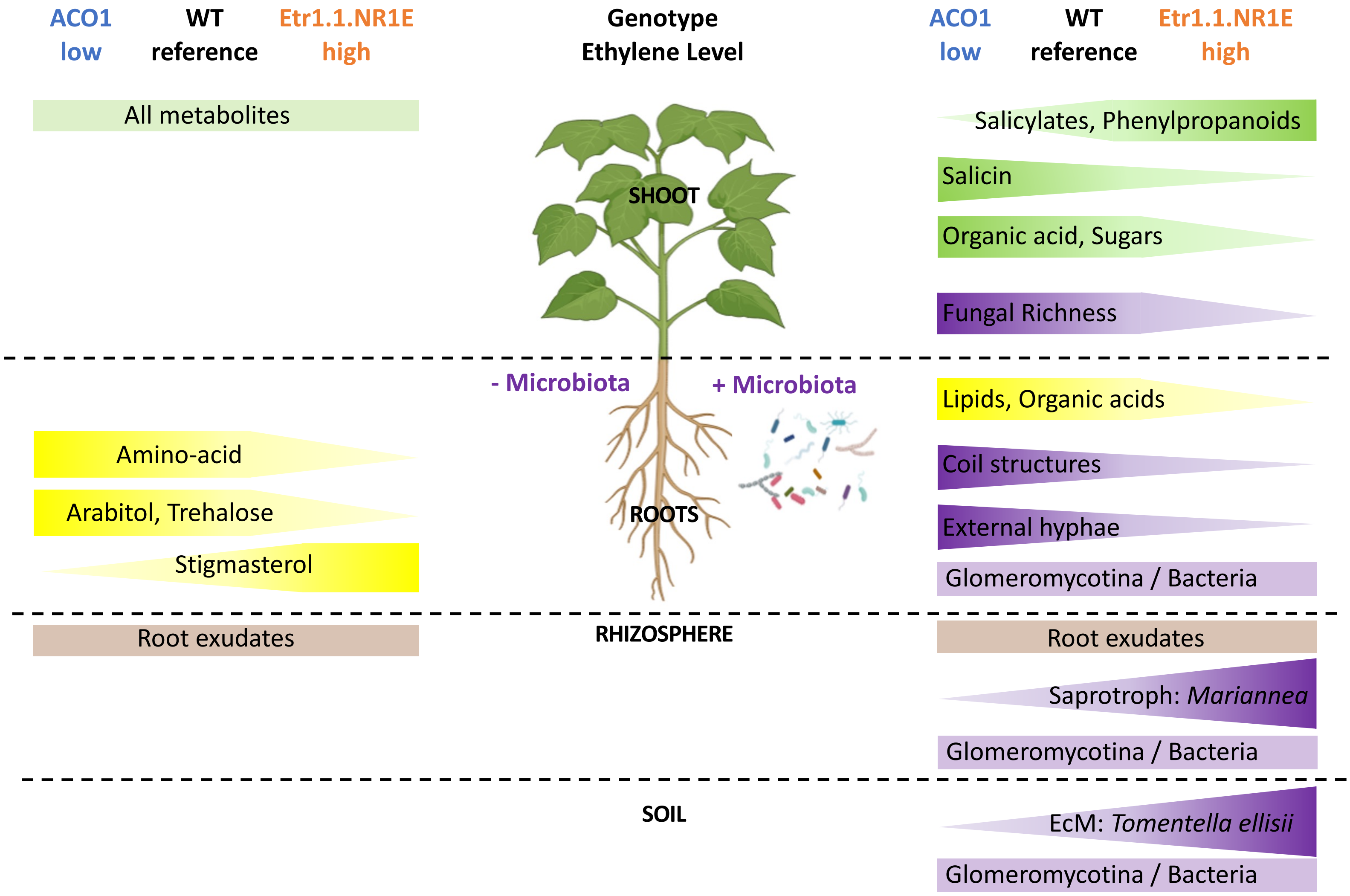
